# Coupling dynamics of 2D Notch-Delta signalling

**DOI:** 10.1101/2022.12.27.521688

**Authors:** Francisco Berkemeier, Karen Page

**Affiliations:** Department of Pathology, University of Cambridge, UK; Department of Mathematics and IPLS, University College London, UK

**Keywords:** Notch, signalling, stability, protrusions

## Abstract

Understanding pattern formation driven by cell-cell interactions has been a significant theme in cellular biology for many years. In particular, due to their implications within many biological contexts, lateral-inhibition mechanisms present in the Notch-Delta signalling pathway led to an extensive discussion between biologists and mathematicians. Deterministic and stochastic models have been developed as a consequence of this discussion, some of which address long-range signalling by considering cell protrusions reaching non-neighbouring cells. The dynamics of such signalling systems reveal intricate properties of the coupling terms involved in these models. In this work, we examine the benefits and limitations of new and existing models of cell signalling and differentiation in a variety of contexts. Using linear and weakly nonlinear stability analyses, we find that pattern selection relies on nonlinear effects that are not covered by such analytical methods.

## Introduction

In epithelial tissue, depending on the nature of the contact between neighbouring and non-neighbouring cells, the Notch-Delta signalling pathway leads to fundamentally different patterns (Cohen, 2003; De Joussineau et al., 2003; Hamada et al., 2014). In highly packed epithelial layers, some cells have the ability to create extensions of themselves, developing protrusions that reach non-neighbouring cells and yielding a new and fundamental factor in the signalling dynamics. These basal actin-based filopodia are elongated and oriented in different directions, working as a signalling arm that extends lateral inhibition to second or third neighbour cells (Hunter et al., 2019; Ramírez-Weber and Kornberg, 1999; Gradilla and Guerrero, 2013; Kornberg and Roy, 2014; Sherer and Mothes, 2008).

In recent years, long-range signalling via filopodia has been shown to significantly impact the distribution and sparse patterning of sensory organ precursor (SOP) cells in the fly notum (Cohen et al., 2010a,b; Hadjivasiliou et al., 2016). In other work, spatiotemporal patterns of spinal neuron differentiation were revealed to be mediated by basal protrusions (Hadjivasiliou et al., 2019). In contrast to the frequently observed salt-and-pepper patterns caused by short-range signalling, cell protrusions result in sparser SOP cell patterning.

The stochastic nature of these biological systems crucially affects patterning. For example, noise arising from dynamic protrusions has been shown to have a significant role in pattern refinement when studying the organisation of bristles on the *Drosophila* notum (Cohen et al., 2010a). A cellular automaton model of cell-cell signalling revealed that rule-dependent structured noise also triggers refined and biased patterning (Cohen et al., 2010b), hinting at the self-organising nature of such systems. Intrinsic noise, driven by stochastic gene expression, has been studied via the Chemical Langevin Equation (Gillespie, 2000) and shown to directly affect juxtacrine-based pattern formation (Rudge and Burrage, 2008).

In addition to constructing realistic long-range signalling models capable of numerically describing long-range patterning, it is also important to develop appropriate analytical tools to understand model behaviour. Linear stability analysis (LSA) has revealed critical and inherent characteristics of lateral inhibition models (Collier et al., 1996; Webb and Owen, 2004; Zakirov et al., 2021; Turing, 1952). Biased and long-range signalling was also studied in Vasilopoulos and Painter (2016), where weight-based coupling functions were considered for several one-dimensional signalling systems. We aim to partially extend this work by studying the twodimensional hexagonal array under specific signalling weights.

We define a model of long-range Notch-Delta signalling, which is a relative weight-based extension of the original Collier model (Collier et al., 1996), and name it the *ϵ*-Collier model. The main idea behind implementing filopodia signalling into the original Collier model is the introduction of a weighting parameter *ϵ* that relatively weights juxtacrine and long-range signalling contributions, creating a complex non-local signalling network. Under different filopodia behaviour and lifespan assumptions, one can explore the robustness of the extended Collier model via LSA, providing a general framework for analysing autonomous systems, as well as one and two-dimensional arrays of signalling cells, as detailed in Supplementary Note 1 (SN1). Furthermore, with such a parameterised model, we aim to investigate the limits on cell coupling sufficient to obtain long-range patterns. In parallel, we explore the effects of stochastic filopodia dynamics and intrinsic noise on patterning. Finally, we expand some of the techniques from LSA to describe a general framework for weakly nonlinear stability analysis (WNSA) of coupled and decoupled dynamical systems.

## Main methods

### Lateral inhibition

We consider a periodic *N* × *M* hexagonal lattice (hexagonal torus), where each cell has 6 neighbour cells. For a given cell, we assume juxtacrine signalling occurs with all 6 of its neighbours via the usual Collier model (Collier et al., 1996). Here, the authors used experimental data to build an ODE model of the feedback loop between two adjacent cells induced by Notch signalling (lateral inhibition). The model consists of a system of coupled ODEs per cell. Denoting by *n*_*i*_ and *d*_*i*_ the levels of Notch and Delta activity in cell *i*, we have the following system

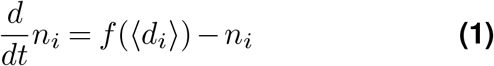

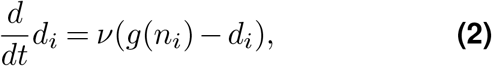

where *f, g* : [0, ∞)→ [0, ∞) are continuous increasing and decreasing functions, often taken to be Hill functions

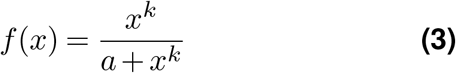

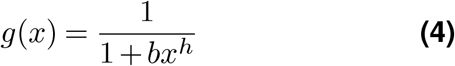

for *x* ≥ 0 and *h, k*≥ 1. *r*_*t*_ ≡ 1/*a* and *b* are the trans-interactions strength and ligand inhibition strength parameters, respectively^1^. *ν* > 0 is the ratio between Notch and Delta decay rates, determining the strength of decay. Finally, ⟨*d*_*i*_⟩ is the average level of Delta activity in the cells adjacent to cell *i*, that is,

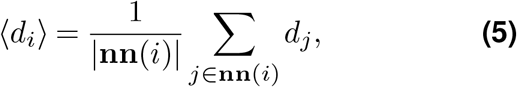

where the sum is taken over the nearest neighbours **nn**(*i*) of cell *i* and |**nn**(*i*)| is the total number of neighbours.

In general, depending on the hexagonal lattice orientation, either *N* or *M* must be even to ensure periodicity. From the previous equations, one can see that the rate of production of Notch activity is an increasing function of the level of Delta activity in neighbouring cells. In contrast, the rate of production of Delta activity is a decreasing function of the level of activated Notch within the same cell. In addition, the production of Notch and Delta activity is balanced by decay.

### Long-range signalling

Besides lateral cell-cell signalling, we also consider the possibility of long-range signalling with respect to non-neighbouring cells. We loosely refer to cell protrusions as the main mechanism for general, isotropic long-range signalling, interchangeably using these terms. A detailed discussion of protrusion dynamics is presented in Mogilner and Rubinstein (2005). For now, our notion of protrusions remains relatively abstract.

There are several ways to implement protrusion-cell signalling. As a first simplification, we assume ⟨*d*_*i*_⟩ is the only term affected by long-range signalling and extend its definition to include non-neighbouring cells that contact cell *i*. Each cell has area 1, and *p*_*ℓ*_ is the maximum protrusion length (or reach) measured from the cell centre. For most of this work, signalling from non-neighbouring cells occurs if a cell is within reach of protrusions. However, we also investigate the cases where ligand density decays with distance and protrusions are stochastic.

In general, we consider the approach suggested in Vasilopoulos and Painter (2016). Here, the authors used a weighting function *ω*(*s, r*) defining the signalling level from a signaller cell *s* to a receiver cell *r. ω* determines which cells are connected through protrusions, defining a connectivity matrix whose entries yield the signalling intensity. In a simplistic protrusion model, all non-zero entries of such a matrix are equal. The weighting function *ω* captures the matrix information, and we may rewrite the interaction term as follows

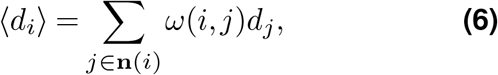

where the sum is made over the neighbours of *i* (**nn**(*i*)) and the non-neighbouring cells that are reached by the protrusions (**np**(*i*)) of cell *i*. Such array of indexes is defined as **n**(*i*) = **nn**(*i*) ∪ **np**(*i*). The further assumption that each cell has a finite amount of active ligand to distribute at any given time point results in the following restriction

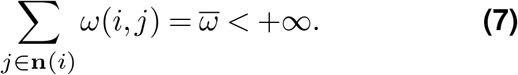

Although there is some freedom in the interpretation of 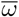, we assume 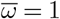 for simplicity.

### The *ϵ*-Collier model

Our model, hereafter named the *ϵ*-Collier model, extends the mathematical systems in Collier et al. (1996) by considering the inclusion of long-range signalling via protrusions balanced by the relative weighting factor *ϵ* ∈ [0, 1].

We begin by weighting each signalling contribution, juxtacrine (*ω*_*J*_) and protrusion-based (*ω*_*P*_), by the factor *ϵ*, to define the combined weighting function

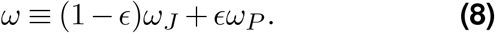

Eq. (6) and Eq. (8) define the *ϵ*-Collier model, considering protrusions of relative signalling intensity *ϵ*. Naturally, Eq. (8) is only interesting when *ω*_*J*_ and *ω*_*P*_ are restricted to **nn**(*i*) and **np**(*i*), respectively. For example, the case *ϵ* = 0 and *ω*_*J*_ (*i, j*) = *χ***nn**(*i*)(*j*)/6, where *χ***nn**(*i*) is the indicator (or characteristic) function of the set **nn**(*i*), corresponds to the original Collier model (Collier et al., 1996).

### Coupling dynamics

We perform a linear stability analysis to understand the criteria for pattern formation driven by Notch-Delta signalling. This is a useful tool to not only identify the regions of the parameter space for which spontaneous patterning of SOP cells occurs but also to determine the typical spacing between Delta-expressing cells, often called the characteristic length of the pattern or pattern wavelength (Turing, 1952; Webb and Owen, 2004; Collier et al., 1996). Our analysis closely follows the methods outlined in Collier et al. (1996); Vasilopoulos and Painter (2016); Formosa-Jordan and Ibañes (2009) and Murray (2001), for the two-dimensional hexagonal array, and is based on the framework presented in SN1. Eq. (1)-Eq. (2) possesses a single positive homogeneous steady state (*n*^*^, *d*^*^). At this state, we have *f* (*g*(*n*^*^)) = *n*^*^ and *g*(*n*^*^) = *d*^*^, which is unique because *f* (*g*(*n*)) is monotonically decreasing for all *n*≥ 0. Then, for small perturbations *ñ*_*i*_ = *n*_*i*_ −*n*^*^ and 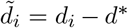, linearisation leads to

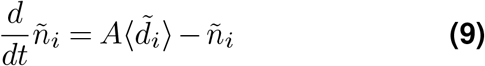

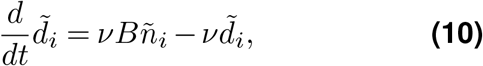

where *A* = *f*^*′*^(*g*(*n*^*^)) is the signal trans-activation by the ligand and *B* = *g*^*′*^(*n*^*^) is the ligand inhibition by the signal. For a *N*× *M* periodic hexagonal lattice, with 1 ≤*j* ≤*N* and 1 ≤*k*≤ *M*, the perturbations can be written as a discrete Fourier series

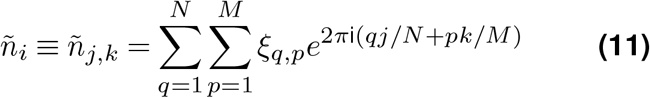

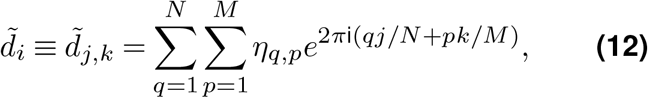

where two subindexes have been used to refer to the spatial position of cell *i* within the two-dimensional hexagonal lattice. For 1≤ *q*≤ *N* and 1≤ *p* ≤*M*, the inverted transform is

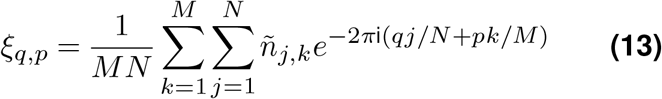

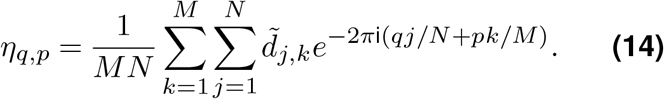

Finally, applying this change of variables to Eq. (9) and Eq. (10) leads to the following system of two ordinary differential equations (ODEs)

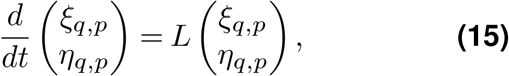

where matrix *L* is a specification of matrix 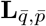 in Eq. (S67), defined as

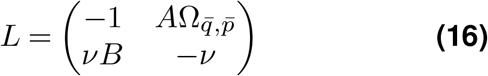

and 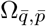 is the function that takes into account the spatial coupling terms of Eq. (9) and Eq. (10) within the hexagonal lattice (in this case, 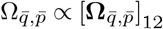, defined by Eq. (S66)). We have then turned Eq. (1)-Eq. (2) into a system of constant-coefficient linear differential equations described by Eq. (15), which has a straightforward family of solutions. For now, however, we focus on the coupling function 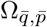, which holds the main mechanisms behind the dynamics of juxtacrine and long-range signalling in our system.

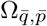 varies according to the weighting function *ω*. Here, 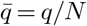 and 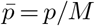 define the discrete wavenumbers (Fourier modes) and thus solutions for 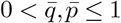 correspond to patterned solutions with corresponding pattern wavelenghts 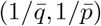. We assume that connections between cells depend only on their relative positions in the lattice (Figure 1b) and therefore, for a sender cell *s* and a receiver cell *r*, we set *ω*(*s*_*j,k*_, *r*_*j*_*′,k′*) ≡ *ω*(*j*^*′*^− *j, k*^*′*^− *k*) = *ω*(Δ*j*, Δ*k*). Hence, 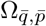 is, in general, given by

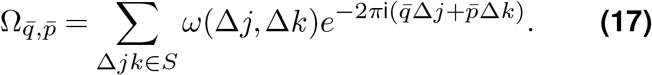

**Figure 1.**
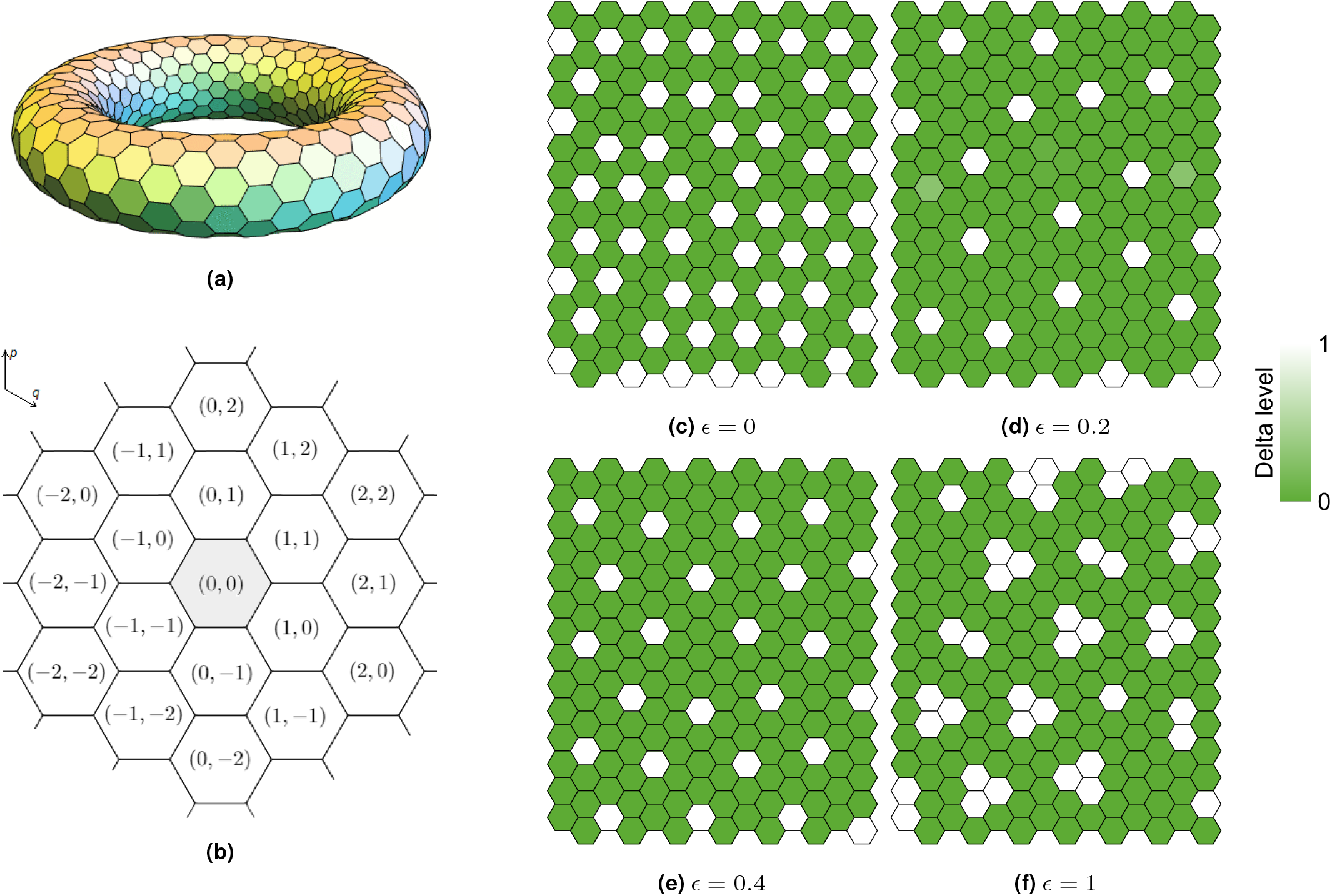
Long-range Notch-Delta signalling on hexagonal lattices. **(a)** Hexagonal torus. Periodic hexagonal lattices can be seen as hexagonal tori. **(b)** Hexagonal lattice main directions (*q, p* axes) and cell position indexation relative to a focal cell (0, 0). Different cell labelling schemes yield equivalent formulations of 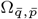. **(c-f)** Notch-Delta patterns on a 14 × 14 periodic lattice for varying *ϵ*. SOP cells (white, high Delta, low Notch) contrast with non-SOP cells (green, low Delta, high Notch). Here, *a* = 0.01, *b* = 100, *h* = *k* = 6 and *ν* = 1. Initial conditions *n*_*i*_(0) and *d*_*i*_(0) have arbitrary values between 0 and 0.1.

Now, if we assume connections are symmetric, i.e, *ω*(Δ*j*, Δ*k*) = *ω*(−Δ*j*, −Δ*k*), we have, by Example 1.1 in SN1,

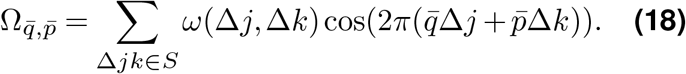

The diagonalisation of *L* leads to the temporal eigenvalues

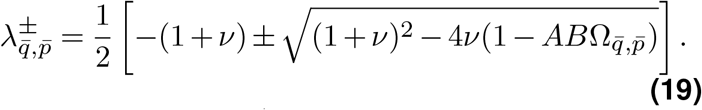

Then, since 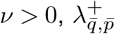 is a positive real number if and only if 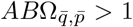. *A* and *B* are the slopes of the feedback functions *f* and *g* at the homogeneous steady state and |*AB*| is defined as the feedback strength. If |*AB*| = 0, then the homogeneous solution is linearly stable, 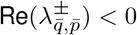, and thus no periodic pattern is expected to emerge. On the other hand, the feedback strength has to be sufficiently high for patterns to arise, that is, |*AB*| *>* |1/Ω_min_|, where Ω_min_ denotes the minimum of the real function 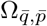, so that 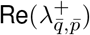 is maximal. With *A >* 0, *B <* 0 and assuming Ω_min_ *<* 0, we expect patterned solutions provided

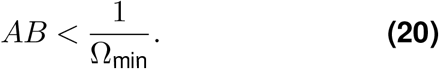

In particular, this feedback is controlled by the tuple (*a, b, h, k*) as follows

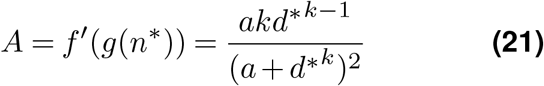

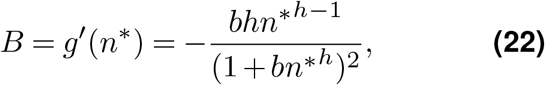

where, again, (*n*^*^, *d*^*^) is a homogeneous steady state, *r*_*t*_ = 1/*a* is the trans-interactions strength and *b* is the ligand inhibition strength. The homogeneous fixed point (*n*^*^, *d*^*^) can be found by setting ⟨*d*_*i*_⟩ = *d*_*i*_ and finding the intersection of *n*_*i*_ = *f* (*d*_*i*_) and *d*_*i*_ = *g*(*n*_*i*_). Assuming for convenience *h* = *k*, this can be rewritten as

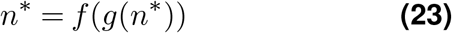

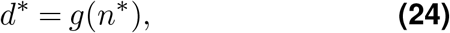

which can be numerically solved for each triple (*h, r*_*t*_, *b*) in the parameter space. Such a solution, together with Eq. (20), defines the discrete Turing spaces consisting of *r*_*t*_-*b* parameter regions there spontaneous patterns occur. Outside such regions, pattern formation is not expected.

We now explore different weighting functions to capture the effects of juxtacrine signalling and protrusions, and discuss what features of *ω* affect Ω_min_. We recall that *ω* determines the family of systems Eq. (1)-Eq. (2) via the weighting dynamics defined by Eq. (6) and Eq. (8).

For a given cell on a hexagonal lattice, we denote the closest ring of order *k* ∈ ℕ_0_ by *R*_*k*_, such that *R*_0_ is the cell itself, *R*_1_ are its 6 neighbouring cells, *R*_2_ is the ring of 12 second-neighbour cells, and so forth. Notice that |*R*_*k*_| = 6*k* (*k* > 0). We further expand the definition of *S* in SN1 by defining *S*_*k*_ as the relative index set of cells in *R*_*k*_ (according to Figure 1b), that is,

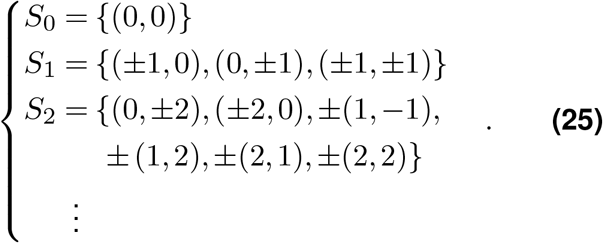

This notation will be used throughout this work. Notice that such a definition can be ambiguous in different contexts, as discussed in Remark 1.2 (SN1).

## Results

### Juxtacrine signalling and simplistic protrusions

For juxtacrine signalling on a hexagonal lattice, without protrusions, we set

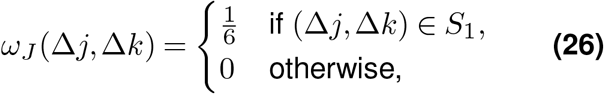

so that

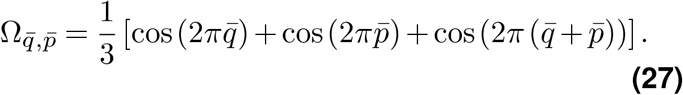

Notice that 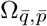 takes discrete values within the interval [−0.5, 1]. The modes that minimise Eq. (27) are those for which *M* and *N* are multiples of 3, thus 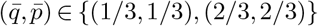 and a pattern with wave-length 3 along the main directions of the hexagonal lattice emerges, provided *AB* < −2. In general, and depending on the initial conditions, such patterns may yield 1, 2 or 3 different cell types, as discussed in more detail below.

Considering protrusions, we first look at the more straightforward case where only the first ring of 12 non-neighbouring cells, *R*_2_, is reached by protrusions. Here, signalling is weighted laterally by Eq. (26) and on *R*_2_ by

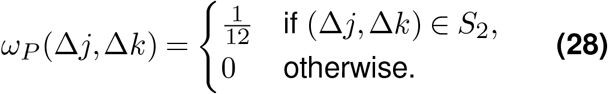

Figures 1c-1f show the observable patterns for different values of *ϵ*, with initial conditions near the homogeneous steady state. Notice that the limit case *ϵ* = 1 has the extreme feature of no juxtacrine signalling, hence the small clusters of Delta-expressing cells in Figure 1f. Even for small values of *ϵ*, sparse patterns are evident. We may then weight each signalling contribution with a factor *ϵ >* 0 and define the combined weighting function *ω* = (1 − *ϵ*)*ω*_*J*_ + *ϵω*_*P*_. Using this leads to

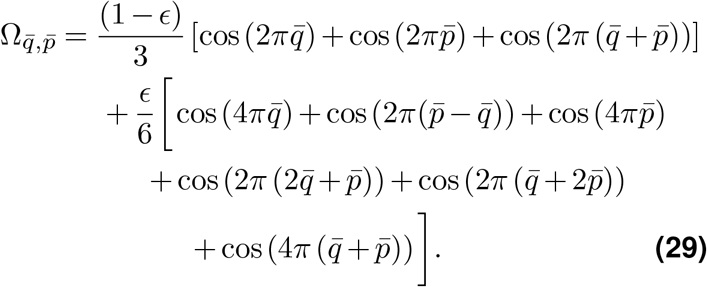

In this case, minimising 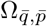 is trickier and therefore we consider a numerical approach. For different values of the long-range signalling strength *ϵ*, Figure 2a shows the change of 1/ |Ω_min_| for increasing values of *ϵ*. Notice that Ω_max_ = 1 for all *ϵ*. Equal juxtacrine-protrusion weighting occurs when *ϵ* = 2/3≃ (Ω_min_(2/3) −0.24). For each *ϵ*, the number of modes varies, as seen in Figure 2b. Notice that 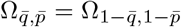 and, in fact, 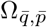 is symmetric with respect to the planes 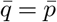 and 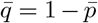 for all *ϵ*. An interesting observation is that at around *ϵ* = 0.4 there are a total of 8 minimising modes, contrasting to the single pair of modes for *ϵ* < 0.4 and the 6 distinct modes for *ϵ* > 0.4 (Figures 2d-2f, Video SV1). The bifurcation observed in Figure 2b at *ϵ* = 0.4 is predicted independently of the Hill functions, and can be mathematically shown by solving, for *ϵ*,

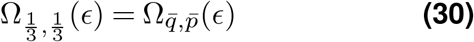

and a minimising pair 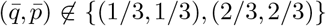 (see SN1 for details). Figures 2g-2i show some of the simulations for corresponding values of *ϵ*.

**Figure 2.**
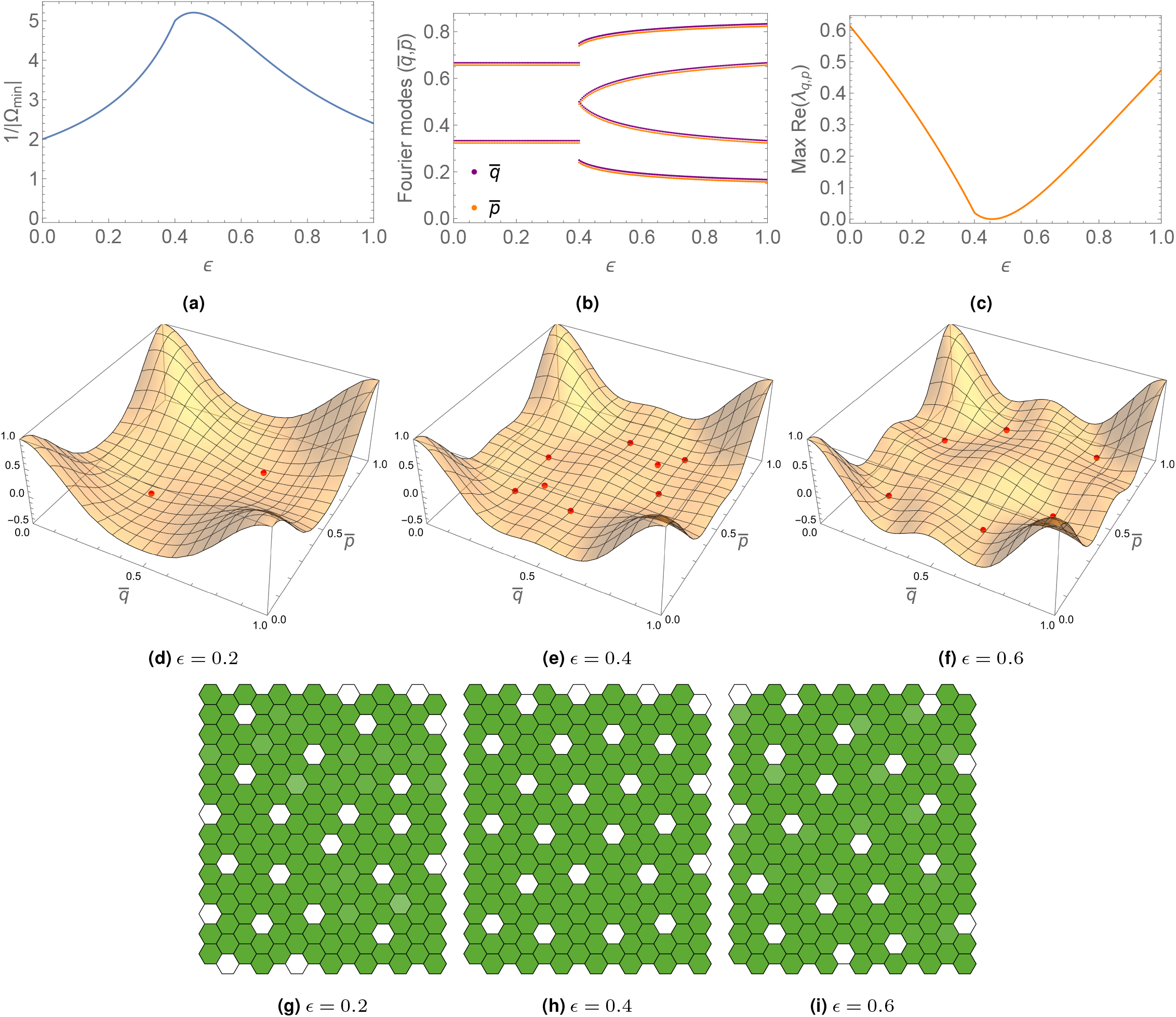
Coupling dynamics of simplistic protrusions as functions of *ϵ*. **(a)** 1/|Ω_min_ |. In this case, 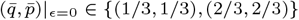 and max_*ϵ*_ Ω_min_ ≃ −0.192. **(b)** Plot of the fastest growing modes 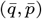. 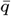 (purple) and 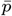 (orange, artificially shifted) have identical plots. **(c)** Maximum value of 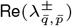 as function of *ϵ*, with *AB* = 1/(max_*ϵ*_ Ω_min_) ≃ −5.207. **(d-f)** Plot of 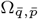 and respective minimising modes (in red). **(g-i)** Simulations on a 14 × 14 lattice for different values of *ϵ*. Here, *h* = *k* = 6. Other parameters are given in Table 1. *n*_*i*_(0), *d*_*i*_(0) ∈ [0, 0.1].

As discussed before, the critical wave numbers maximise the real part of the temporal eigenvectors. Equivalently, Figure 2c shows 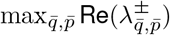 as a function of the relative weight parameter *ϵ*, corresponding to the critical *AB* = 1/(max_*ϵ*_ Ω_min_) ≃ −5.207. Here, 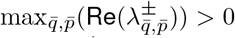 for all *ϵ*, and thus patterns are expected to emerge with the maximising wavelength modes. As suggested by Eq. (21)-Eq. (24), we may go a step further and work out the specific parameter regions for which |*AB*| yields pattern formation. The phase diagrams (Turing spaces) in Figure 3a (Video SV2) represent the regions in the *r*_*t*_-*b* plane such that *AB* < 1/Ω_min_, or more specifically,

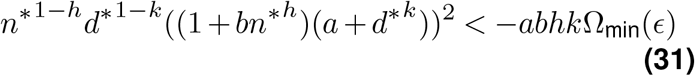

for different values of Ω_min_ and corresponding *ϵ*. The *ϵ*-Collier model is quite robust with respect to the pair (*r*_*t*_, *b*), corresponding to the trans-interactions strength and ligand inhibition strength parameters, respectively. Increasing *ϵ* from zero initially reduces the size of the discrete Turing space, in which patterning occurs, followed by an increase after intermediate values of *ϵ* (Ω_min_(*ϵ*) has a maximiser at *ϵ* ≃ 0.455), which is in accordance with the monotonicity change of 1/ |Ω_min_| (Figure 2a). Note that Turing spaces for different values of *ϵ* are strictly contained sets, via Eq. (31). In the region (*r*_*t*_, *b*) ∈ × [10−1, 108] [100, 102], patterns emerge for any *ϵ*, since 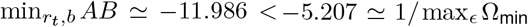 (there are always non-empty Turing spaces).

**Figure 3.**
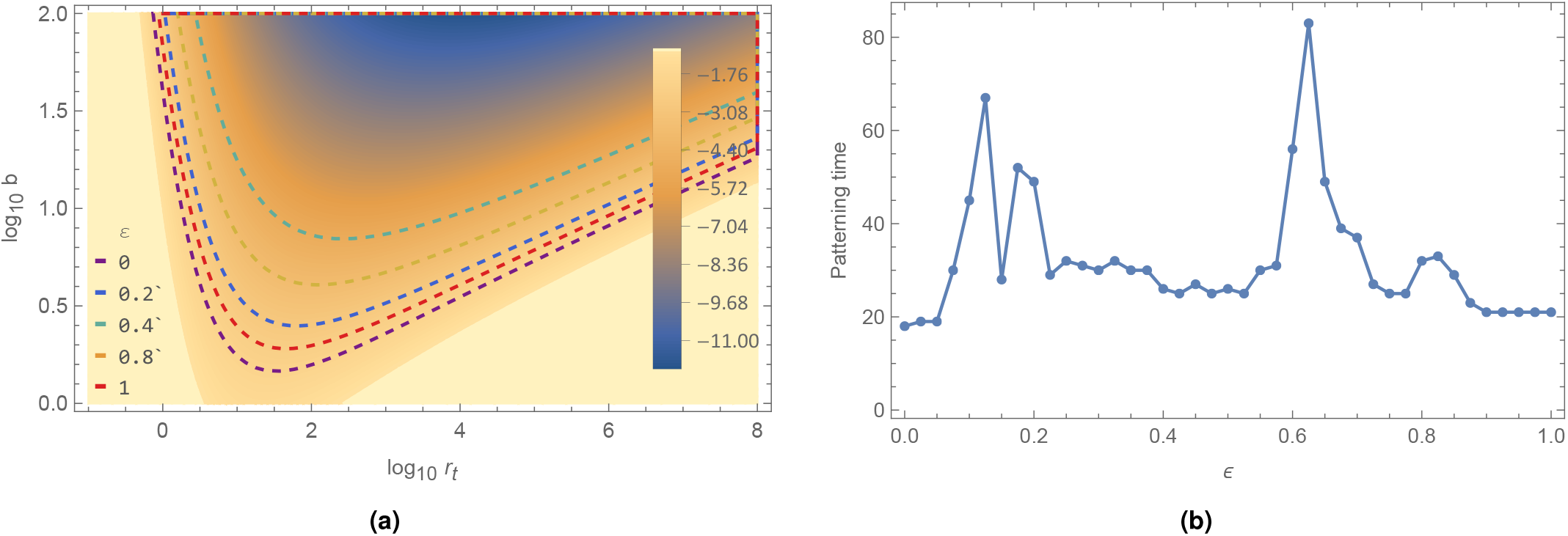
Phase diagrams and patterning speed. **(a)** Log-log contour plot of *AB* as a function of *r*_*t*_, *b*. The regions delimited by the dashed lines indicate the Turing spaces where spontaneous pattern formation occurs (*AB* < 1/Ω_min_(*ϵ*)), for each *ϵ* ∈ {0, 0.2, 0.4, 0.7, 1}. Here, *h* = *k* = 6. In this region, 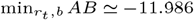 at (*r*_*t*_, *b*) ≃ (10^4.041^, 10^2^). Purple line corresponds to *AB* = −2 (*ϵ* = 0). **(b)** Patterning time for different values of *ϵ*. Blue dots indicate the mean patterning time for each *ϵ*. Here, 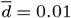. Parameter values are given in Table 1. *n*_*i*_(0), *d*_*i*_(0) ∈ [0, 0.1].

Patterning speed is also affected by relative weighting, as seen in Figure 3b. We define the patterning time as the first instant *t* for which

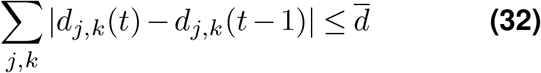

for some threshold 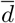. Fixing 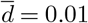, we see that, patterning slows down for different ranges of *ϵ*, resulting in slowing spikes around 0.05 < ϵ < 0.2 and 0.6 < *ϵ* < 0.7. Simulations were done over tissues with random initial conditions, for each *ϵ*.

### Multiple cell types

For a narrow range of *ϵ* values, Eq. (1)-Eq. (2) converges into alternative stable solutions that include more than two cell types (based on stable Delta activity levels). This effect is observable at both ends of the *ϵ* spectrum, defining thresholds of pattern selection.

To two decimal places, for *ϵ* ≃ 0.039, 6× 6 and 14 ×14 periodic lattices yield approximately 3 and 5 different cell types, respectively (Figures 4a-4b). For *ϵ*≃ 0.85, we get approximately 3 and 4 cell types, respectively (Figures 4c-4d). Whether some of these solutions eventually converge to others, reducing the number of distinct cell types, is not known. The definition of a cell type is also somewhat ambiguous, but we believe this effect to be noteworthy nonetheless.

**Figure 4.**
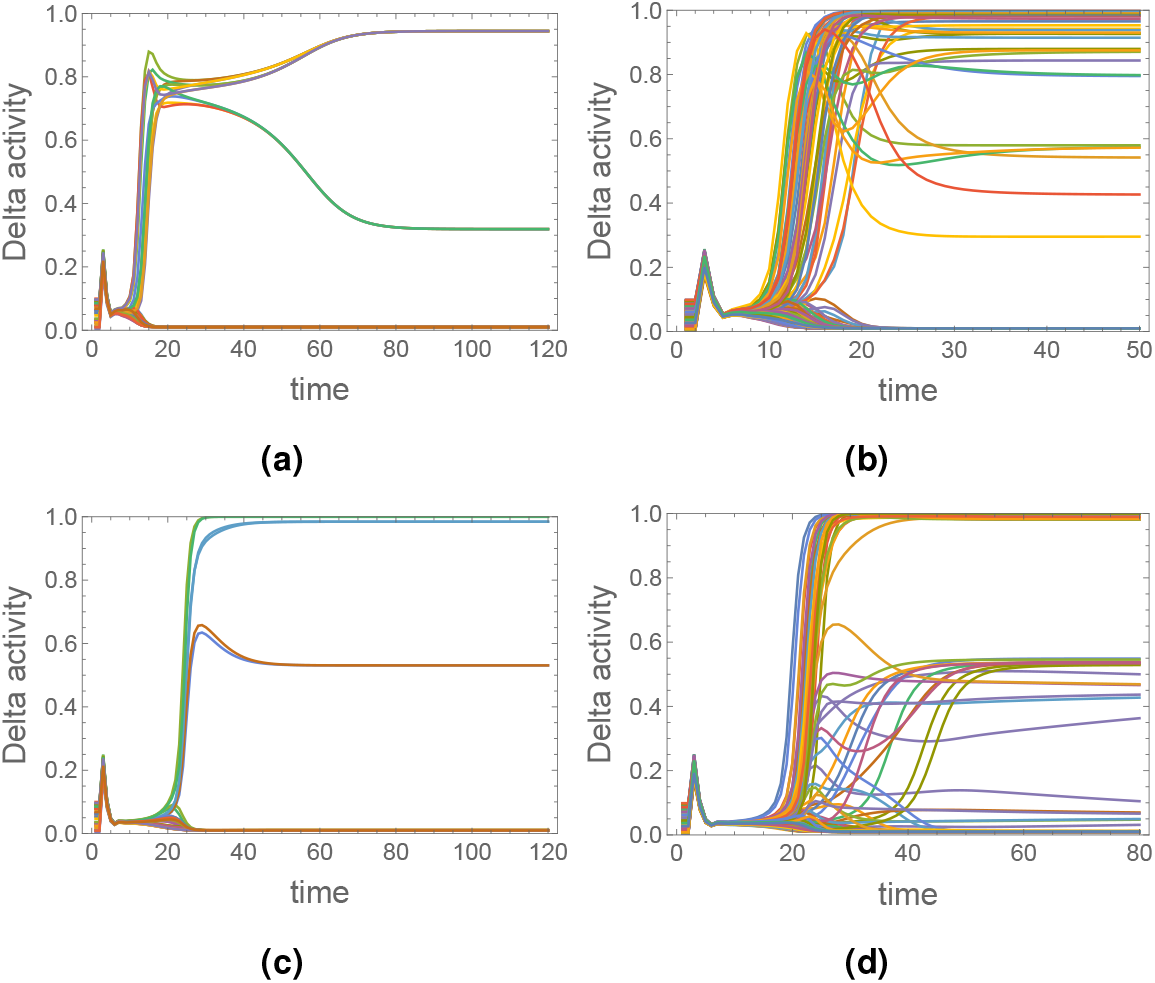
Dynamics at critical *ϵ* values. Delta activity dynamics on 6 × 6 **(a**,**c)** and 14 × 14 **(b**,**d)** periodic lattice, for **(a-b)** *ϵ* ≃ 0.039, and **(c-d)** *ϵ* ≃ 0.85.

### Long and oriented protrusions

A possible first extension is to consider the effects of longer or oriented protrusions. Examples of applications regarding this type of long-range signalling can be found in the emergence of spotted patterns similar to the ones observed in the skin of *pearl danio* fish and striped *zebrafish*-type patterns (Hamada et al., 2014; Eom et al., 2015; Kondo et al., 2021). In Vasilopoulos and Painter (2016), a general weighting function was considered to account for protrusion length and orientation. Here, we adapt such a framework by focusing only on the protrusion weighting component 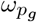, given by

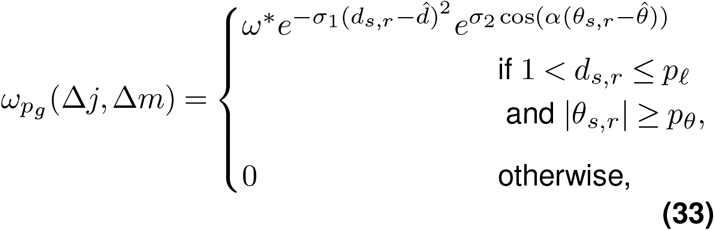

where *d*_*s,r*_ and *θ*_*s,r*_ ∈ (−*π, π*] are defined as the relative distance and the angular bearing between the signalling and receiving cells, respectively. *σ*_1_, *σ*_2_, 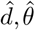 and *α* are parameters that control the shape and form of the weighting function, and *ω*^*^ is the normalising coefficient implicitly defined by Eq. (7). Furthermore, *p*_*ℓ*_ is the maximum protrusion length and *p*_*θ*_ is an angle bound.

We assume *d*_*s,r*_ is the same for each cell in *R*_*k*_, *k*≥ 2, and thus we may rewrite it, using our previous index notation, as a function of (Δ*j*, Δ*m*)

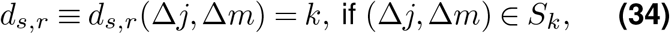

which yields 1 < *k* ≤ *p*_*ℓ*_. Definition Eq. (34) is equivalent to the hexagonal-Manhattan distance *d*_*H*_ defined in Remark 1.2. Notice also that, with *p*_*ℓ*_ = 2, *p*_*θ*_ = 0, *σ*_1,2_ = 0 and *ω*^*^ = 1/12, we recover the *R*_2_ weighting function *ω*_*P*_ given by Eq. (29). We set, in this case, 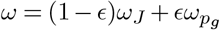.

We focus only on the case of longer protrusions and thus we impose radial symmetry by taking *σ*_2_ = 0, *p*_*θ*_ = 0 (cases with *σ*_2_ > 0 lead to axial and polarised signalling systems, as discussed in Vasilopoulos and Painter (2016)). Intuitively, *σ*_1_ represents the strength of ligand density decay with distance. In the following, we assume long-range signalling strength to decrease as a function of *d*_*s,r*_ and therefore take 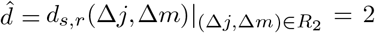 and *σ*_1_ > 0. Hence Eq. (33) simplifies to

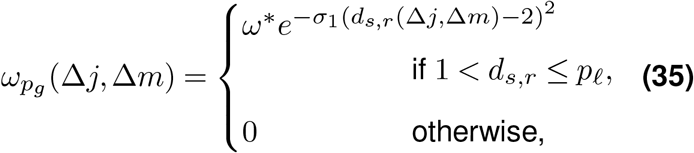

where, from Eq. (7),

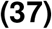

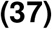

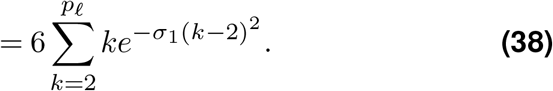

Under the assumption that protrusions may reach up to *R*_4_ (*p*_*ℓ*_ = 4), we have that, as an example, 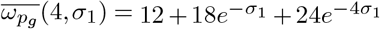. For different values of *σ*_1_, Figure 5d shows the minimal feedback strength required for patterning, derived from 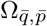. The case *σ*_1_ = 0 yields equal *R*_*k*_ (2 ≤ *k* ≤ 4) weighting and thus 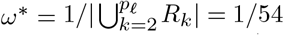, in this case (Figures 5a-5c). As *σ*_1_ → ∞, *ω*^*^ → 1/12 and we recover the dynamics for *R*_2_ protrusions (Figure 5e). Figures 6a-6c show simulations for different values of *p*_*ℓ*_. Surprisingly, the pattern wavelengths for *p*_*ℓ*_ = 4 are not correctly predicted by LSA, as seen by comparing the minimisers of Figure 5c with the simulation in Figure 6c. For this value of *p*_*ℓ*_, the minimising modes remain unchanged for a wider range of *ϵ*.

**Figure 5.**
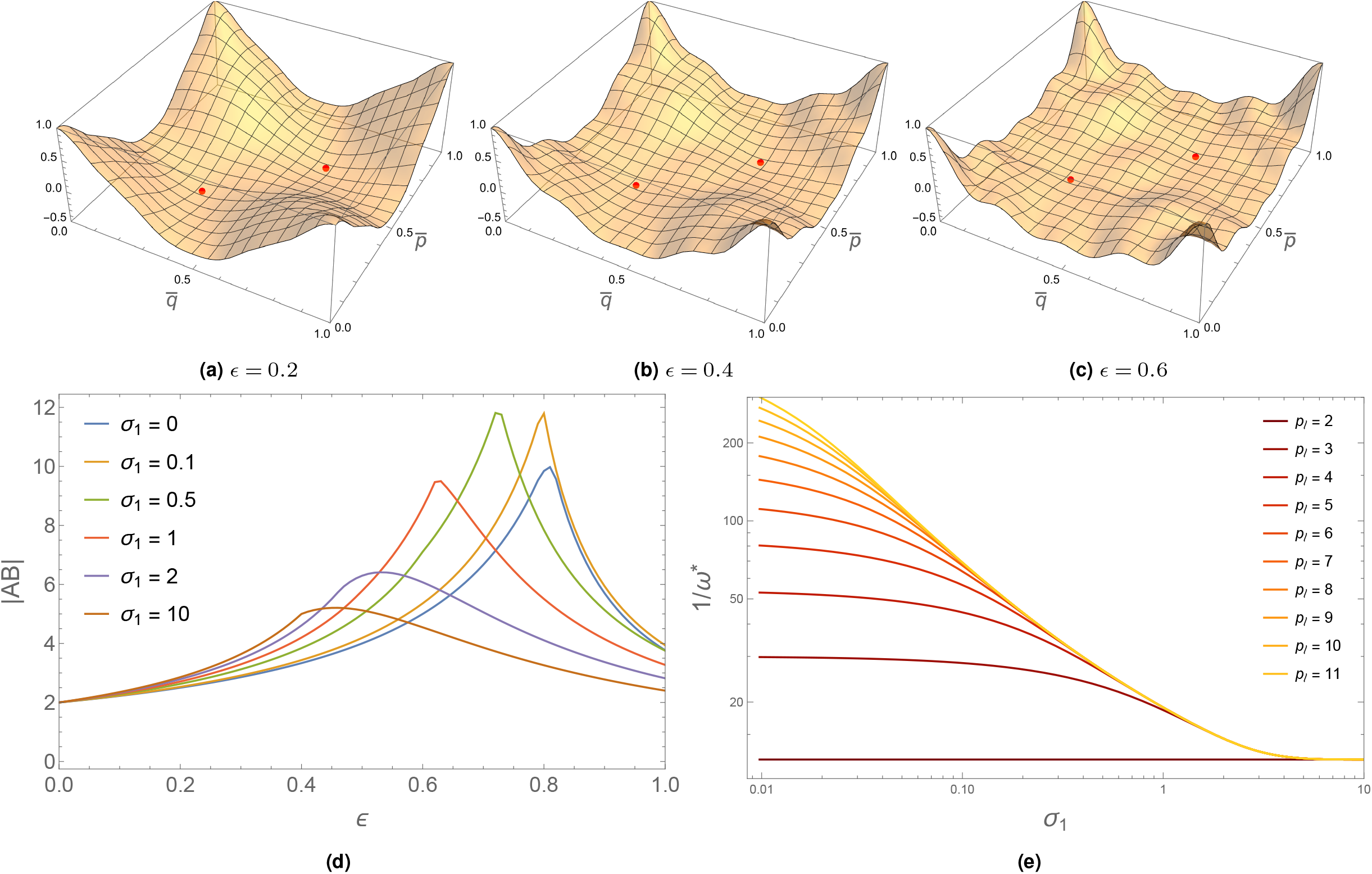
Coupling dynamics of long protrusions as functions of *ϵ*. **(a-c)** Plot of 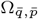 and respective minimising modes (in red) with *σ*_1_ = 0. Wavier plots are observable due to the more complex nature of the weighting function, with *p*_*ℓ*_ = 4. **(d)** Critical |*AB*| for different values of *σ*_1_, given by |1/Ω_min_|, with *p*_*ℓ*_ = 4. **(e)** 1*/ω*^*^ as a function of *p*_*ℓ*_ and *σ*_1_. *ω*^*^ → 1/12 as *σ*_1_ → *∞*. Other parameters are given in Table 1.

**Figure 6.**
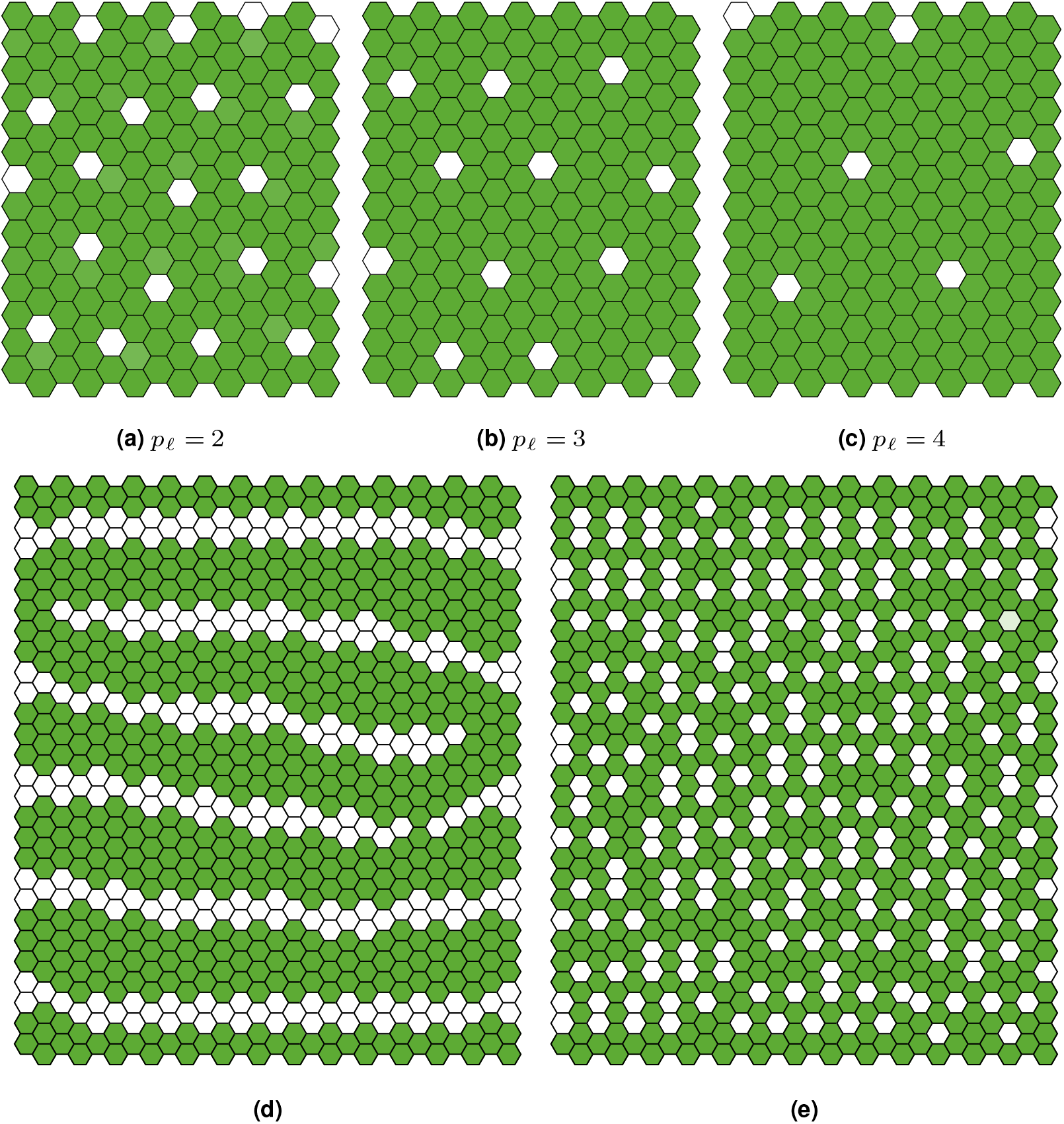
Patterning with long and oriented protrusions. **(a-c)** Simulations on a 14 ×14 lattice for different values of *p*_*ℓ*_. Here, *ϵ* = 0.6 and *h* = *k* = 6. **(d-e)** Bounded protrusions may lead to other patterns, relevant to other applications. Here, *ϵ* = 1. Parameter values for all simulations are shown in Table 1 in Supplementary Note 3.

Considering bounded protrusions significantly alters the coupling function and symmetry may be broken. Interesting pattern may arise in this case when *ϵ* = 1, specially regarding the emergence of clustering effects or *zebrafish*-type patterns (Binshtok and Sprinzak, 2018; Kondo et al., 2021; Moreira and Deutsch, 2005) (Figures 6d-6e).

### Stochastic protrusions

One way of generalising the weighting function *ω* is to consider some level of randomness in protrusion-cell signalling. Previous studies have suggested that pattern regularity and refinement can be greatly improved by considering dynamic lifespan-based protrusions (Cohen et al., 2010a). Here, we extend such an approach to the *ϵ*-Collier model on the *R*_2_ ring.

Depending on the protrusion type and level of biological detail, different stochastic models may be implemented. For example, in the case of the eukaryotic flagellum (Marshall and Rosenbaum, 2001) and stereocilia (Narayanan et al., 2015), the length evolution can be studied using a master equation with length-dependent rates of protrusion attachment and detachment. For such systems, the length fluctuations can be mapped onto an Ornstein-Uhlenbeck process. In another case, Pilus, which is a bacterial protrusion, keeps elongating and retracting with velocities in between pauses (Koch et al., 2021), and so the length dynamics can be described by a three-state Markov process. Further details on the physics of filopodial protrusions can be found in Mogilner and Rubinstein (2005); Patra and Chowdhury (2020) and an extensive discussion on length control of long cell protrusions of various types was presented in Patra et al. (2022).

Here, however, we assume isotropic protrusions and consider dynamic binding and unbinding of filopodia to non-neighbouring cells throughout the simulation. The lifespan of protrusions is determined by birth and death rates, *p*_*b*_ and *p*_*d*_, respectively. These correspond to the attachment and detachment rates of protrusions to non-neighbouring cells. Within a short time interval Δ*t*, a link is formed between a cell and one of its second neighbours with probability *p*_*b*_Δ*t*. Such a link is destroyed with probability *p*_*d*_Δ*t*. This leads to the formulation of a continuous-time *telegraph process* (Kac, 1974; Bena et al., 2002; Kolesnik and Ratanov, 2013; López and Ratanov, 2014) with rates *p*_*b*_ and *p*_*d*_. This process is also known as a *dichotomic* or two-valued Markov process. In the following, we first present well-known results on telegraph processes, followed by the application to our case.

From stochastic theory, the general telegraph process is defined as a memoryless continuous-time stochastic process that has two distinct values. If the two possible values that a random variable *X*(*t*) can take are *x*_1_ and *x*_2_, then the process can be described by the following master equations

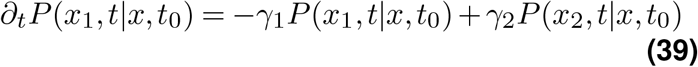

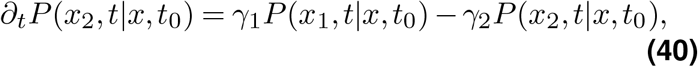

where *γ*_1_ is the transition rate from state *x*_1_ to state *x*_2_ and *γ*_2_ is the transition rate from state *x*_2_ to state *x*_1_. In matrix form, Eq. (39)-Eq. (40) can be rewritten as

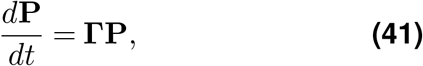

where **P** = (*P* (*x*_1_, *t*|*x, t*_0_), *P* (*x*_2_, *t*|*x, t*_0_))^*T*^ and

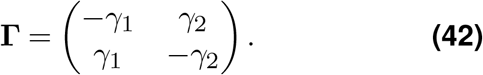

To find the general solution of Eq. (41), we begin by determining the eigenvalues and eigenvectors of matrix **Γ**, which are given by *λ*_1_ = 0, *λ*_2_ = −(*γ*_1_ + *γ*_2_) and

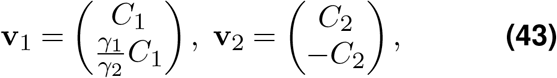

respectively, with *C*_1_ and *C*_2_ depending on the initial conditions. Hence, the solution is then given by

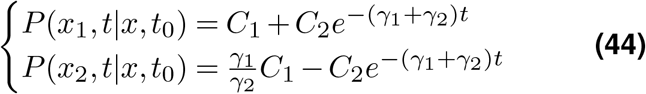

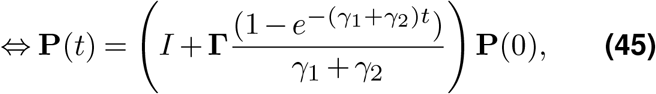

where **P**(0) = [*C*_1_ + *C*_2_, (*γ*_1_*/γ*_2_)*C*_1_ − *C*_2_]^*T*^ in the case *t*_0_ ≡ 0, without loss of generality. With initial conditions given by

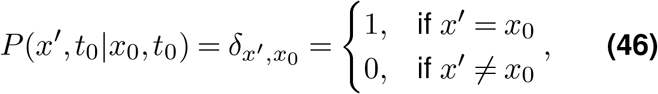

where *δ*_*ij*_ is the Kronecker delta, the solution in the compact form can be given by

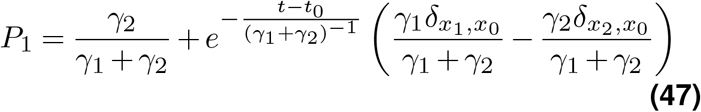

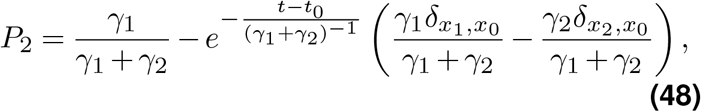

where *P*_1_ = *P* (*x*_1_, *t* | *x*_0_, *t*_0_) and *P*_2_ = *P* (*x*_2_, *t*|*x*_0_, *t*_0_). We are now interested in studying the asymptotic dynamics of the telegraph process, approximating its discrete realisations to a Bernoulli process, given a suitable condition on the realisation timescales. With Δ*t*≡*t*−*t*_0_ ≫ (*γ*_1_ + *γ*_2_)^−1^, the solution approaches a stationary distribution **P**_*s*_ given by

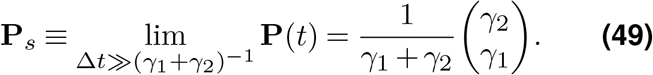

The time-dependent ensemble average satisfies

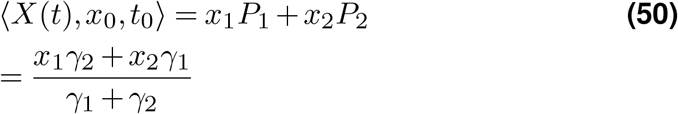

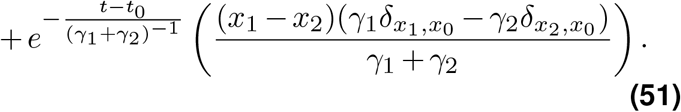

Hence, the stationary average is given by

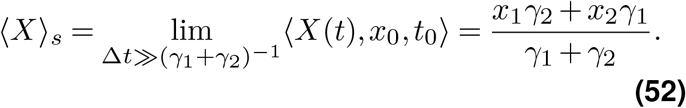

In our case, we have *γ*_1_ = *p*_*b*_, *γ*_2_ = *p*_*d*_, *x*_1_ = 0 and *x*_2_ = 1. Hence, in the limit where Δ*t* ≫ (*p*_*b*_ + *p*_*d*_)^−1^, the probability of finding a protrusion is *p*_*b*_/(*p*_*b*_ + *p*_*d*_). We may then treat such a process as a Bernoulli process with probability *p*_*b*_/(*p*_*b*_ + *p*_*d*_). In other words, if the timescale at which we make the observation is longer than the inverse of the event rates, we may expect the process to be memoryless every time we observe, describing a Bernoulli process.

In the following, we assume that neighbouring *R*_1_ cells are always linked, with weight (1 − *ϵ*)/6, and *R*_2_ cells are linked with stochastic weight 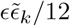. *ϵ* is now weighted by independent and identically distributed (i.i.d.) random variables

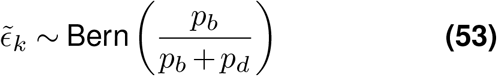

for each *k* in *R*_2_. At each time step, the stochastic coupling term is then given by

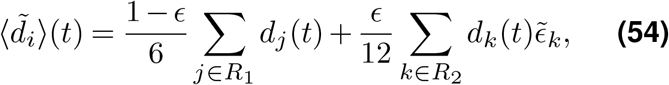

where the second term is a sum of weighted i.i.d. Bernoulli distributions. Note that 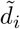 here should not be confused with the homogeneous state perturbation introduced in other sections. One of the key aspects of having dynamic protrusions is the possibility of pattern refinement over time. As suggested in Cohen et al. (2010a), we define the coefficient of variation, *ζ*_*V*_, of a pattern as the ratio between the standard deviation and mean of the distances from each SOP cell to its 6 closest SOP cells. This coefficient yields a measure of the global order of the emergent pattern, which we then track for different values of (*p*_*b*_, *p*_*d*_), as seen in Figure 7a. The case (*p*_*b*_, *p*_*d*_) = (10, 3.5) is particularly interesting as the pattern converges to ideal cell packing (*ζ*_*V*_ = 0) at around *t* = 390 (Figure 7e).

**Figure 7.**
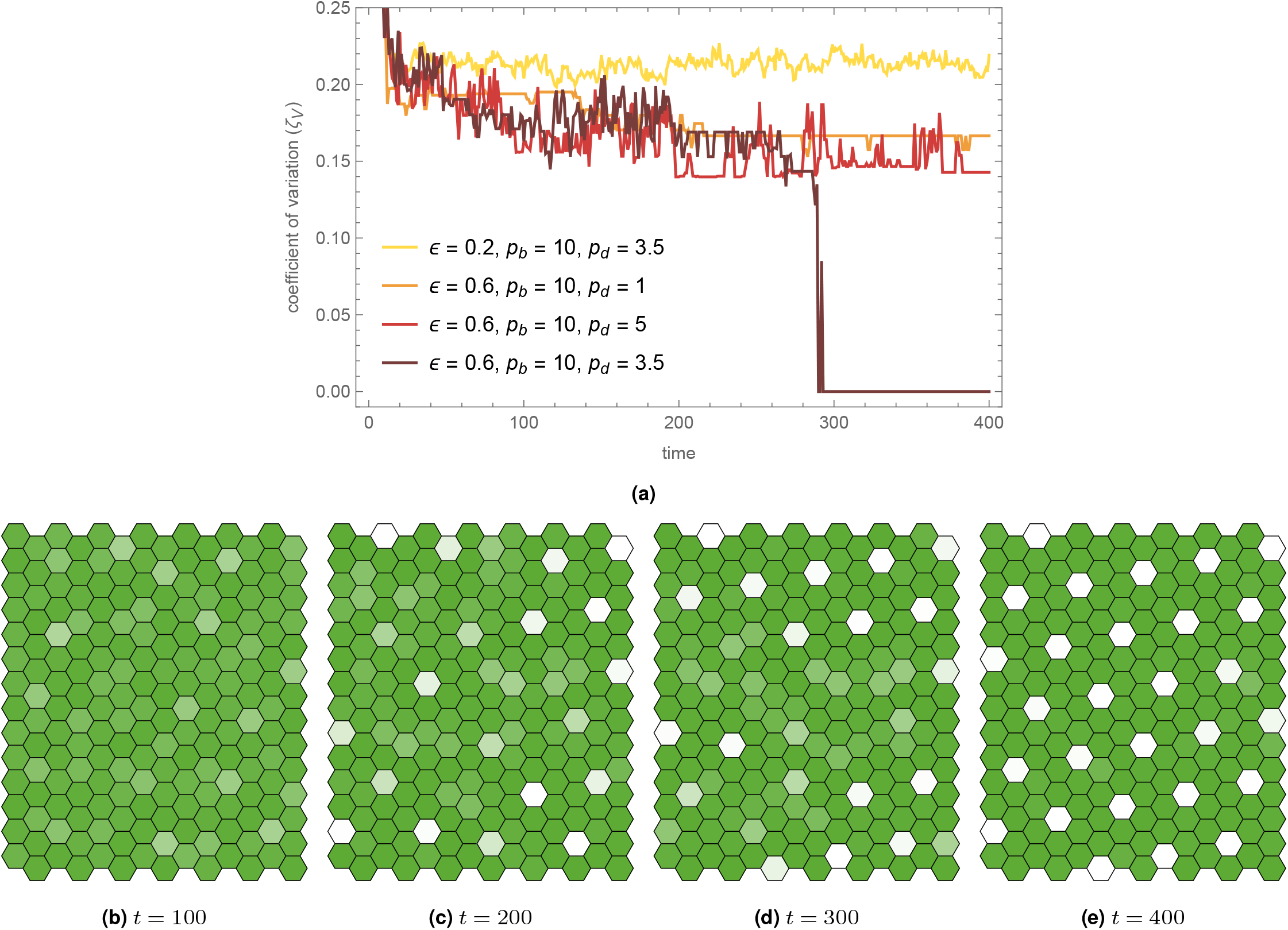
Dynamics of stochastic protrusions. **(a)** Stochastic behaviour of the coefficient of variation *ζ*_*V*_ in spacing between nearest Delta-expressing cells. **(b-e)** Pattern refinement may take a long time to stabilise. Once reaching the refined state at *t* = 400, the pattern hardly changes. The final pattern is also stable under no protrusions. Here, *ϵ* = 0.6, *p*_*b*_ = 10 and *p*_*d*_ = 3.5. Other parameter values are given in Table 1.

Some patterns only stabilise once optimal packing is attained, depending on the tissue dimensions. For a periodic tissue whose dimensions are multiples of 14, which guarantees optimal *R*_2_-signal sparse patterning is possible (notice that 14 × 7 would also work), once the coefficient of variation is minimised, patterning does stabilise. In many cases, given enough time, the stable *R*_3_ pattern is obtained after gradual refinement determined by *p*_*b*_ and *p*_*d*_. In a perfectly refined pattern, one should expect *ζ*_*V*_ = 0, which means each SOP cell is surrounded in *R*_3_ by 6 other equally spaced SOP cells (Figure 7e, Video SV3).

It should be noted that the simulations shown in Figure 7 are all isolated examples corresponding to single realisations. Although the purpose of this study is to identify *p*_*b*_ and *p*_*d*_ such that pattern refinement is achieved, the complex relation between such rates to guarantee convergence to the refined pattern may be hinted at by a thorough stochastic analysis, which is beyond the scope of this work.

Interestingly, if we look at the extreme case of sudden removal of protrusions (from the refined state), we see that such a state is stable under purely juxtacrine lateral inhibition (Figure 7e). This is similar to taking *ϵ* = 0 after pattern stabilisation. Patterns of such wavelength contrast with the ones predicted by LSA, but they do not contradict pattern selection under consideration of non-linear terms (Collier et al., 1996), as discussed below. Considering different approaches to noise-driven protrusions might help in better understanding the role of stochastic effects in patterning and refinement. For instance, avoiding the Bernoulli approximation on the Markov-type protrusion dynamics could hint at a more realistic description of filopodium behaviour and consequently pattern formation. Noise-mediated filopodium reach and orientation have been studied in Cohen et al. (2010a). Cellular automaton models have also been used to explain sparse and more complex patterns (Cohen et al., 2010b). Dichotomous noise has also been applied in Langevin dynamics, in a broader scenario (Barik et al., 2006). Hence, a natural alternative to this source of noise, is to study the role of intrinsic noise, driven by Langevin dynamics, which we discuss next.

### Intrinsic noise

Different types of noise have been considered in other works, specifically of additive and multiplicative nature. The additive noise approach considers noise as a stochastic term that is added to each of the species equations in the signalling system (in our case, two) in the form of independent Wiener processes. The multiplicative noise approach is subject to response functions that are proportional to the response level of the system. In other words, the noise term is multiplied by a factor that scales with the current state of the system. The key difference is that additive noise has fixed magnitude, while multiplicative noise depends on the level of the state variables (Fuliński and Telejko, 1991; Van Kampen, 1992).

In this section, we aim to study the impact of intrinsic noise on statistical fluctuations in SOP cell patterning via long-range signalling as per the *ϵ*-Collier model. We also study the patterning speed depending on the effective cell volume within the multiplicative noise terms. In case of morphogen-mediated patterning of gene expression, intrinsic noise has proven to affect timescale dynamics of bistable switches (Perez-Carrasco et al., 2016). Stochastic effects were shown to accelerate juxtacrine pattern formation and alternative lateral inhibition models (Wearing et al., 2000) were found to be robust to intrinsic noise (Rudge and Burrage, 2008). Statistical properties of protein concentration in gene-regulated networks were more generally discussed in Thattai and Van Oudenaarden (2001). In order to motivate the investigations of this section, we revise well-known facts about the Chemical Langevin Equation (CLE), a multivariable Itô stochastic differential equation that describes the time evolution of molecular counts of reacting chemical species (Gillespie, 2000). In signalling systems, noise may arise from the stochastic nature of chemical kinetics derived from random motions and collisions of molecules. For *K* molecular species, if **X**(*t*) = (*X*_1_(*t*), …, *X*_*K*_(*t*)), where *X*_*k*_(*t*) is the number of molecules of species *k* in the system, and provided **X**(*t*) is a jump-type Markov process under a propensity-based evolution law, chemical kinetics regarding *M* reaction channels can be captured by the Chemical Master Equation (CME) (Mcquarrie and Gillespie, 1967; Gillespie, 1976, 1992), given by

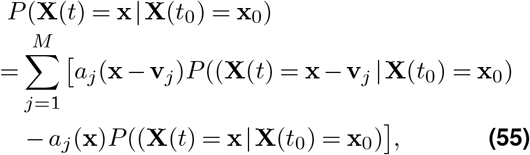

where **v**_*j*_ is the stoichiometric (state-change) vector for reaction *j*. The propensity function *a*_*j*_ is mathematically defined as the product between the specific probability rate constant and the number of distinct combinations of reactant molecules available in the state **X**, for each reaction channel (Oppenheim et al., 1969; Kohlmaier, 1972). The Stochastic Simulation Algorithm (SSA) presented in Gillespie (1977) often accompanies the CME as an exact approach to simulating chemical kinetics. Here, however, we focus on the stochastic differential equation approximation, discussed next.

For well-mixed systems with large numbers of molecules, the species dynamics can be deterministically captured by

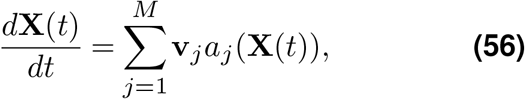

which is often written in terms of the species concentrations. In Gillespie (2000) it was shown that, within “macroscopically infinitesimal” time intervals (so that propensity functions remain approximately constant, yet many reactions are expected to occur), the CME Eq. (55) can be approximated as a chemical Langevin equation (Gillespie, 1996; Turner et al., 2004; Li and Li, 2017), given in its standard white noise form as

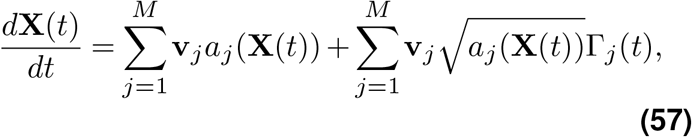

where the Γ_*j*_(*t*) are temporally uncorrelated, statistically independent Gaussian white noises. Specifically, Γ_*j*_(*t*) satisfy ⟨Γ_*j*_(*t*)Γ_*k*_(*t*^*′*^)⟩ = *δ*_*jk*_*δ*(*t* − *t*^*′*^). Here *δ*_*jk*_ is the Kronecker delta, *δ*(*t* − *t*^*′*^) is the Dirac delta.

In our case, we are interested in understanding how noise generated by chemical kinetics influences longrange signalling dynamics. While Notch and Delta levels may have ambiguous definitions (Collier et al., 1996), here we assume they are concentrations, thus requiring an adaptation of Eq. (57) using the effective cell volume *V*, given by the ratio between the average number of species molecules and the maximum values *n*_*i*_ and *d*_*i*_ attain. In some sense, *V* defines the noise intensity in these signalling dynamics. With the change of variables (*n*_*i*_, *d*_*i*_) = **X**(*t*)*/V*, Eq. (57) becomes, for each cell *i*,

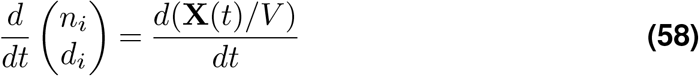

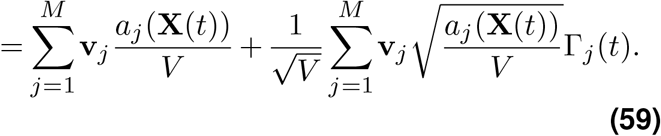

We may then write the stochastic version of Eq. (1)-Eq. (2)

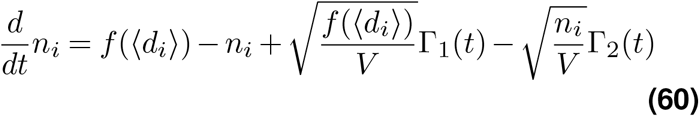

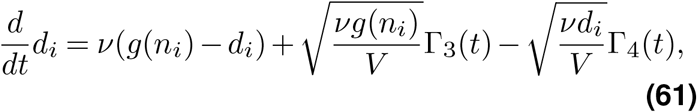

where we fixed *M* = 4, *a*_1_ = *f* (⟨*d*_*i*_⟩)*V, a*_2_ = *n*_*i*_*V, a*_3_ = *νg*(*n*_*i*_)*V* and *a*_4_ = *νd*_*i*_*V*. Notice that Γ_*j*_ are also uncorrelated for different cells, though we avoid over-notation with index *i*. In our case, *K* = 2 and *M* = 4 for *N* cells. Eq. (57) is known as the standard-form CLE, but it is not the only possible formulation, and other formulations with different numbers of Gaussian noises are conceivable (Schnoerr et al., 2014; Mélykúti et al., 2010). In this section we discuss stochastic simulations based on Eq. (60)-Eq. (61). Note that the deterministic system is recovered when *V* → ∞.

Figure 8 shows the simulation results for different values of *ϵ* and effective volume *V*, within values motivated by the discussion in the previous paragraph. Effective volume values greater than 1000 seem to approximate the deterministic case. Note that the different realisations for the deterministic case (*V* → ∞) correspond to different initial conditions. Patterning time was computed by identifying a threshold percentage of the mean Delta activity of the deterministic steady state solution. Once a stochastic system passed such a threshold, the time was registered. It is interesting to notice that patterning time does not seem to correlate with the effective volume. A clear delay is observable for an intermediate value of *ϵ* (*ϵ* = 0.4), when compared to the deterministic case. However, it is worth noting that simulations were done over a relatively small number of lattices to optimise computational efficiency. More simulations might potentially yield different results.

**Figure 8.**
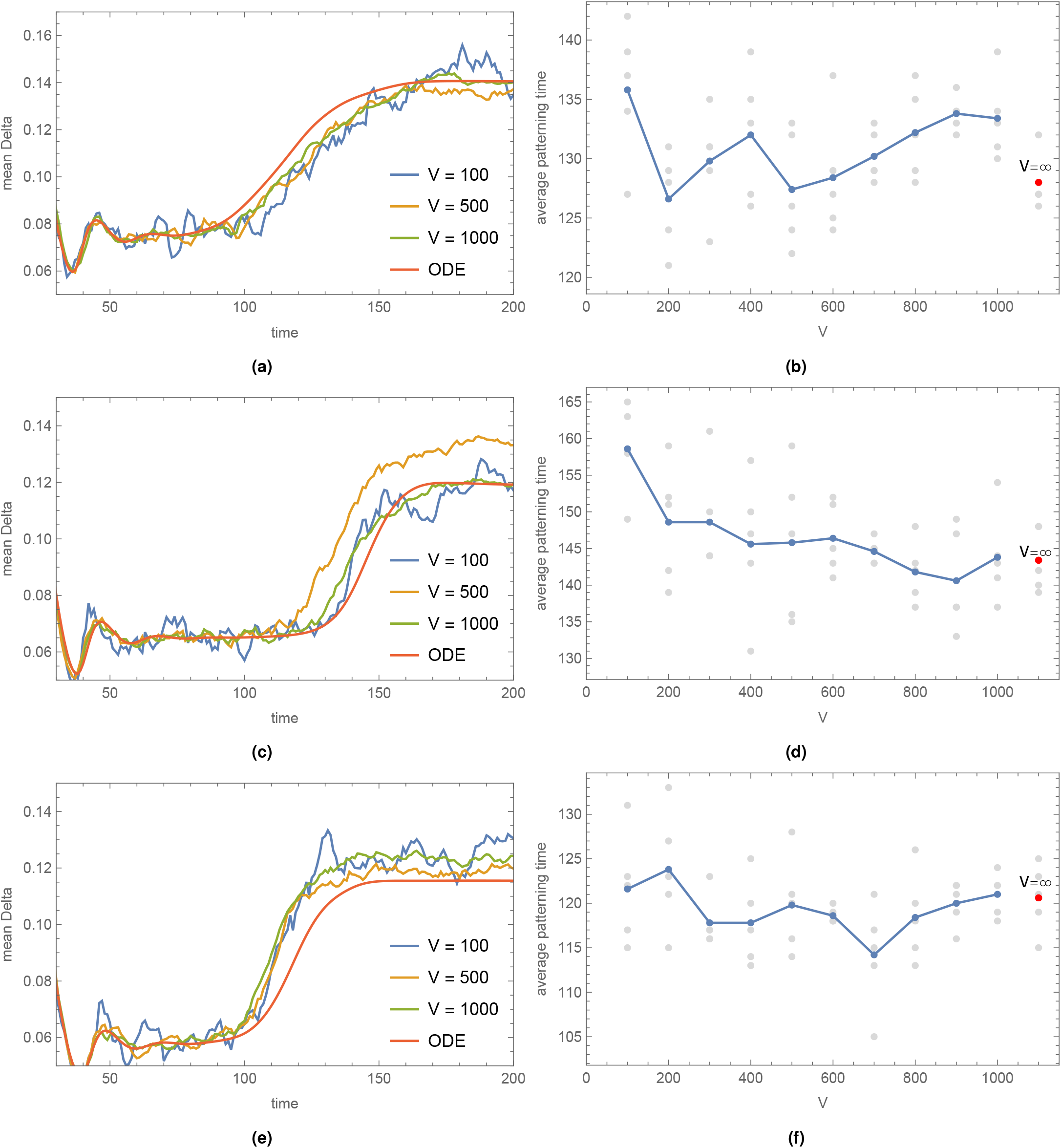
Intrinsic noise dynamics with long-range signalling. Plots on the left show the mean Delta activity dynamics over time. Plots on the right show the patterning time. Blue dots indicate the average over 5 samples (grey) per cell volume, while the red dot corresponds to the limit *V*→ ∞ (deterministic ODE). Simulations were done for different values of *V* and *ϵ* on 14 × 14 periodic lattices. Here, *h* = *k* = 6 and the mean Delta threshold was set to 0.12 (all cells). (a-b) *ϵ* = 0.2. (c-d) *ϵ* = 0.4. (e-f) *ϵ* = 0.6. See Table 1 for details.

### Robustness and pattern selection

We now explore how Fourier analysis describes pattern selection under LSA. Again, we discuss robustness to changes in two of the main parameters in the *ϵ*-Collier model: the Hill function switch parameters, given by the trans-interactions strength *r*_*t*_ = 1/*a* and the ligand inhibition strength *b*. We study the convergence to the desired pattern with long-range signalling for different values of *a* and *b*. Here, we consider protrusions acting on the *R*_2_ ring. In a system with two variables per cell like the linearised system

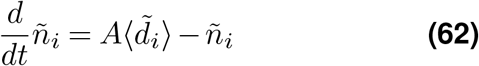

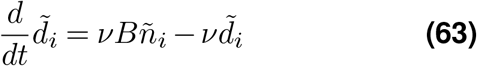

the characteristic polynomial is of second order. As a consequence, each couple 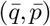 has two eigenvectors and eigenvalues. We then have that the solution of the linearised problem is given by

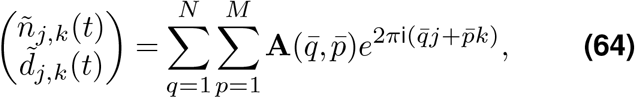

where

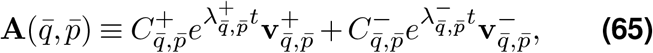

and where 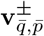 and 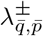 are the eigenvectors and eigenvalues associated to 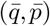, respectively. 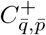 and 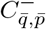 are constants depending on the initial conditions of the problem. In the case that at least one family of modes 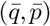 grows exponentially fast, it linearly destabilises the homogeneous solution and this family dominates over the rest, giving rise to a periodic pattern with the 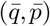-wavenumbers. At such critical modes, we have that the eigenvalue with the largest real part and respective eigenvector are given, as functions of *ϵ*, by

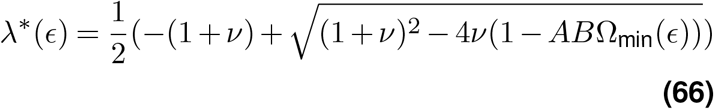

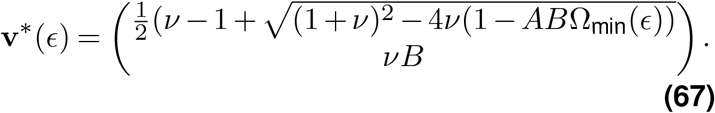

Hence, for large *t*, the dominant pattern is a superposition of modes with periodicity determined by 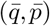 and *ϵ*. Thus, for each family of critical modes 𝒲, the solutions in Eq. (64) asymptotically satisfy

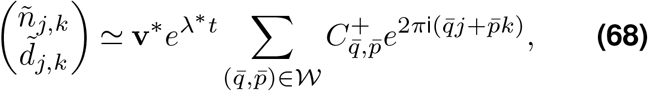

where *λ*\*, **v**\* and 𝒲 are all functions of *ϵ*. It is then clear that the long term behaviour of this solution is dependent on the amplitudes of the Fourier components in Eq. (68), which in turn depend on the initial conditions of the problem. In fact, since 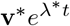 is independent of 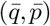 and 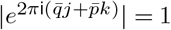, the relative amplitude is given by 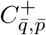, which is implicitly determined by

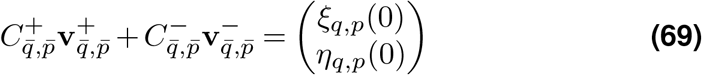

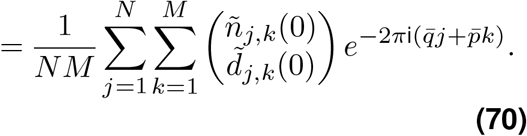

Hence, depending on the choice of the initial conditions and consequently, 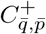, the long term behaviour of the solution could yield different patterns and cell types. Similar to the analysis in Collier et al. (1996), the generic pattern predicted by linear stability analysis might yield more than two cell types, depending on the choices of 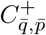.

Numerically, however, cells usually opt for one of two possible fates, (*n, d*) ∈ (0, 1), (1, 0), and therefore nonlinear effects play a role in determining the number of cell types. Consequently, our model is robust because the final pattern of cell differentiation is not affected by the specific form of the Hill functions, as long as the feedback between cells is strong enough, similar to the lateral inhibition case (*ϵ* = 0).

Figure 9 shows the pattern selection with corresponding fastest growing modes for different values of *ϵ* and 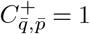. From vector Eq. (68), we simply plot the real part of the normalised sum values of its second term,. -corresponding to Delta activity (determining the opacity of each white cell). When necessary, and to illustrate the nature of the pattern, we provide rational approximations of the real values of the minimising 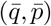wavenumbers.

**Figure 9.**
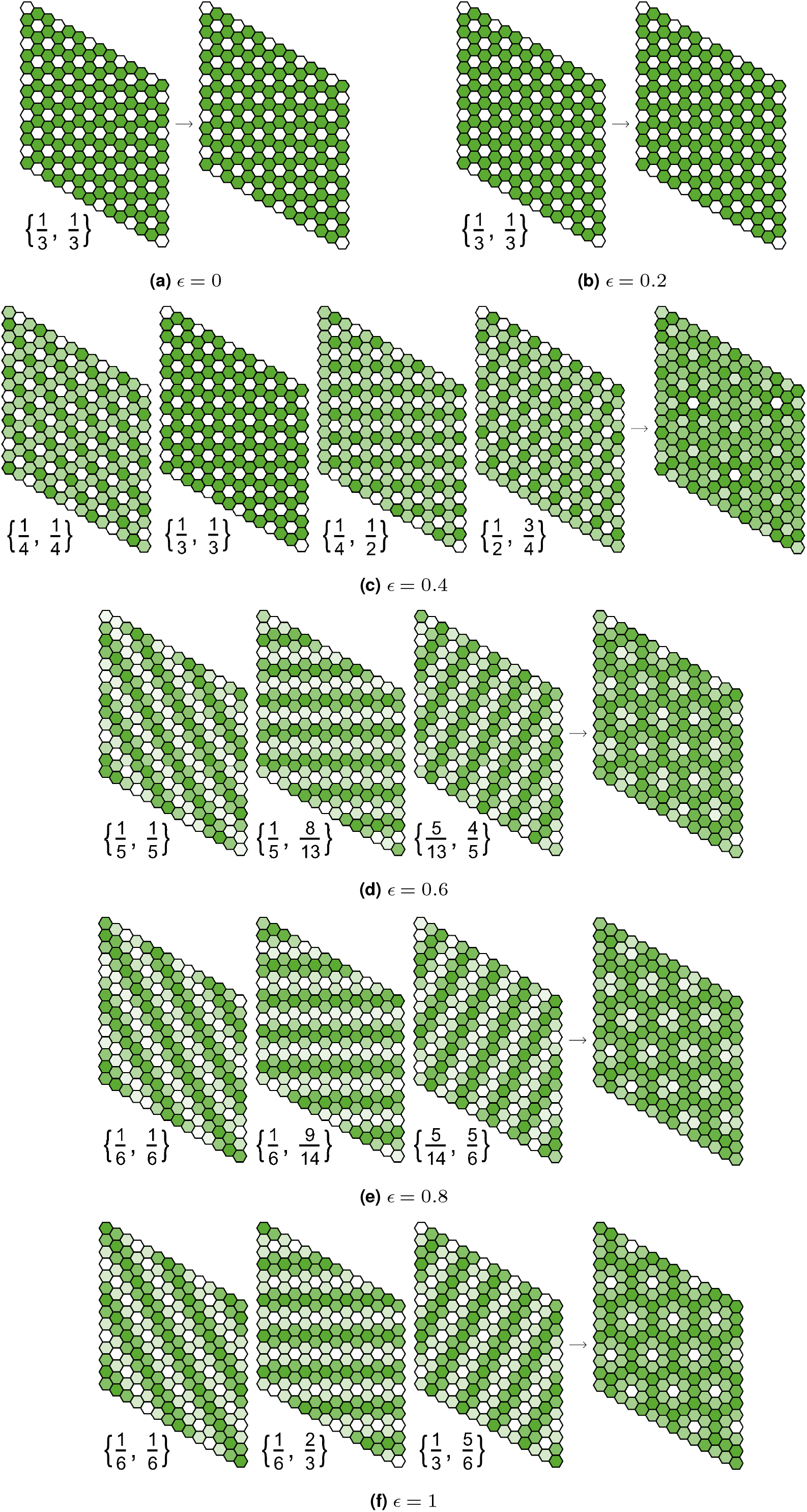
Pattern selection via LSA: asymptotic solutions. Eq. (68) decomposes the asymptotic solution to the linearised system in terms corresponding to different minimizing modes 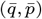. Each term solution is represented on the left of each of the subfigures **(a-f)**, for different values of *ϵ*, while their linearly combined solution is on the right. Due to symmetry, the pattern observed for each single mode 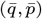 is the same for 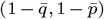 and thus we omit half of the single mode patterns. Here, 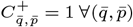. The modes in **(d)** and **(e)** are rational approximations of the real values, as explained in the main text. Other parameter values are given in Table 1.

Figure S5 compares the final patterns from Figure 9 with an SOP cell filtering based on a specific threshold (percentage of steady state solution) and a numerical simulation. In other words, the plots in each middle panel correspond to the selection of cells whose Delta level is above a specific threshold (*d*_*T*_). It is noticeable that LSA fails to predict sparse patterning for a wide range of values of *ϵ*. It is only for *ϵ* ≥ 0.4 that sparser patterns emerge, yet more than two cell types are observable prior to filtering. Figure S6 shows the dependence of cell fates on the choice of 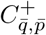 Similar to the conclusions for lateral inhibition in Collier et al. (1996), different choices of constants lead to fundamentally different SOP cell patterns and cell types. Considering imaginary constants, for example, leads to intermediary cell types in the case of 0 ≤ *ϵ* < 0.4 (Figure S6a). Despite partially describing sparse patterning via long-range signalling, LSA does not fully capture the main mechanisms required to explain longer wavelength patterns. A nonlinear approach is then required to extract some more information on the dynamics of such systems and pattern selection.

### Weakly nonlinear stability analysis

Since LSA has failed to predict sparse pattern formation for many values of *ϵ*, alternative types of stability analysis could be explored. In reaction-diffusion systems, weakly nonlinear stability analysis (WNSA) has provided further qualitative conclusions about the effects of nonlinear terms, based on the amplitude dynamics of the Fourier components corresponding to the fastest growing modes (Turing, 1952; Wollkind et al., 1994). In Supplementary Note 2 (SN2), we explore a possible application of such methodologies to patterning derived from long-range signalling. In the following, we take **u**_*i*_ = (*n*_*i*_, *d*_*i*_) and

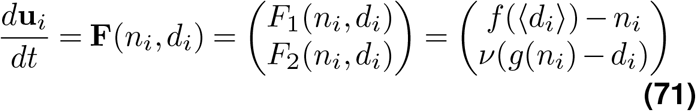

for each cell *i*. We aim to extend the linear approach to consider solutions of Eq. (71) in the harmonic form

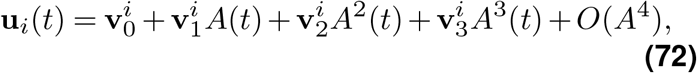

for some constant vectors 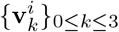, and where *A* satisfies the following Landau equation (Aranson and Kramer, 2002)

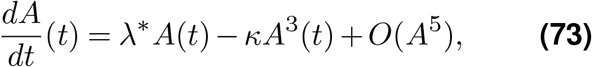

where *λ*\* and *κ* are motivated in SN2. Discrete Fourier transforms may be used to decouple the original system of 2*NM* equations. In the linear case, given exponential-based solutions of the linearised system, the main problem relied on minimising 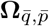 in order to find the fastest growing modes 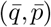. This problem changes and becomes relatively trickier in the case of WNSA due to multiple mathematical obstacles.

The extension of 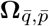 to the weakly nonlinear solution in Eq. (72) is not trivial and, in general, we should not expect a higher-order extension of the decomposition in Eq. (S164) to occur due to cross-derivative terms. However, given the specific shape of our system (see SN2 for details), the higher-order cross-derivatives are all zero. Hence, we may write

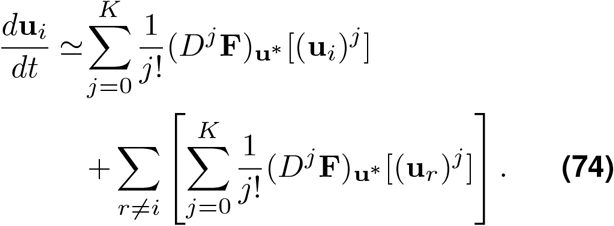

where (*D*^*j*^**F**)_**u**_* is defined in SN2. All that remains now is to find a simplification of the right-hand side term of Eq. (74) so that the decoupling is complete and we may write the entire expression as a function of (*n*_*i*_, *d*_*i*_).

As detailed in SN2, however, such a decoupling seems mathematically unfeasible in the case of translationally invariant Notch-Delta signalling systems, given the complexity generated by the higher-order terms of Eq. (74). Any methodology as systematic as the linear case seems to be out of reach within our framework. Therefore, WNSA is insufficient to describe quantitative dynamics of long-range signalling, without further assumptions, as discussed below.

## Discussion

In this work, we have outlined some of the main tools for analysing a general long-range signalling model. We developed such a model by taking a relative signalling approach. Long-range signalling via filopodia is weighted by a parameter *ϵ*, while the juxtacrine contribution is weighted by 1 − *ϵ*. This constitutes the *ϵ*-Collier model, understood as a long-range extension of the original Collier model (Collier et al., 1996). We found that sparser patterns on periodic hexagonal lattices are robust and tend to emerge even for small values of *ϵ*. Additionally, we discovered that the speed of patterning is dependent on *ϵ*, directly delaying or accelerating differentiation for different ranges of *ϵ*.

To comprehend the linear effects of long-distance signalling, we first employed linear stability analysis for generally coupled and translationally invariant systems, followed by a direct application to the *ϵ*-Collier model of long-range Notch-Delta signalling. We explored various protrusion modelling situations, including shortand long-range protrusions, stochastic protrusions, and intrinsic noise dynamics. While LSA proved to be a useful tool for identifying the fastest-growing modes under Fourier analysis, we also revealed some of its faults in the long-range scenario. In contrast to solely juxtacrine models, LSA failed to predict sparser patterning across a broad range of *ϵ* values, indicating that nonlinear effects play a significant role in sparse pattern selection. Nonetheless, LSA predicted sparse patterns for certain values of *ϵ* given a particular SOP detection threshold. We also examined the system’s response to various assumptions on the effective volume of cells and found that the effective volume appears to have no effect on the patterning time. However, this was performed on small lattices for computational efficiency, and the results may vary for other lattice sizes.

Motivated by such limitations in LSA, we devised a framework for weakly nonlinear stability analysis in order to achieve expanded qualitative conclusions on wavelength selection in Fourier-transformed coupling functions. Specifically, we presented the main methodology behind a potential framework for WNSA of translationally invariant Notch-Delta systems. We expanded well-known reaction-diffusion techniques to coupled spatially discrete systems, such as the *ϵ*-Collier model on a periodic hexagonal lattice, by considering harmonic-based Landau-type solutions. Unfortunately, we found that the decoupling mechanism in translationally invariant systems appears impractical in WNSA.

For the purpose of refining our LSA and WNSA estimates, bifurcation theory may shed light on other critical parts of our analysis. Numerically, one might additionally examine parameters closer to the point at which the homogeneous state becomes linearly unstable (dashed lines in Figure 3a). Here, it is anticipated that the model will behave more linearly; hence, LSA may match the simulation outcomes better. If this does not occur, it may indicate a subcritical bifurcation (as opposed to a supercritical one), and when a subcritical bifurcation occurs, LSA does not necessarily predict the patterning outcome (Crawford, 1991; Stefanou and Alevizos, 2016). In the case of WNSA, we computed higher-order terms around an equilibrium branch. Similarly, if we focus instead on a bifurcation point, we may be able to make progress and solve the mathematical dilemma.

This work provides a framework for understanding pattern formation in relatively general long-range signalling systems, highlighting some of the primary mathematical obstacles in such a theory and indicating possible generalisations and future research avenues.

## Supporting information

SV1-Coupling_function

SV2-Turing_spaces

SV3-Stochastic_protrusions

SV4-Manhattan_patterns

## Computational methods

### Simulations

All simulations were performed using *Interactive Epithelium* (IEp), a *Wolfram Mathematica* tool for hybrid Notch-Delta epithelial signalling and patterning simulations. IEp aims to provide a practical tool for testing parameter robustness while simulating the dynamics of the Notch-Delta signalling pathway in an epithelium. Table 1 in Supplementary Note 3 contains the precise parameter values for the simulation plots shown in this work.

### Code availability

The source code and data that were used to develop the main conclusions and analyses presented in this work are available on the following repository hosted on GitHub: https://github.com/fberkemeier/Notch-DeltaCoupling.git. The relevant video simulations can also be found in this repository. Previous versions are available upon request. For any comments/suggestions, as well as copyright issues, please contact.

## ACKNOWLEDGEMENTS

We thank Dr. Mohit Dalwadi at University College London and Dr. Nick Monk at the University of Sheffield and AIMS Ghana for the comments that greatly improved the manuscript. We thank Dr. Giulia Paci, Prof. Yanlan Mao and Prof. Buzz Baum for helpful discussions.

## Footnotes

1 The quantity *a*^1/*k*^ is the neighbour Delta activity level necessary for half-maximal Notch activation, while *b*−^1/*h*^ is the Notch activity level necessary for half-maximal Delta inhibition.

