## Supplementary Information for "Coupling dynamics of 2D Notch-Delta signalling"

### Supplementary Note 1: Linear stability analysis

This section provides an introductory and fairly general approach to linear stability analysis (LSA). First, we discuss the main methods used in LSA and present some examples and applications, leading to the main study case: the Notch-Delta signalling pathway, in the main text.

#### Main methods

Consider the $n$-dimensional dynamical system, represented by the vector $\mathbf{x}(t) = (x_1(t), x_2(t), ..., x_n(t))^T$ and described by the ODE system

$$\dot{\mathbf{x}} = \mathbf{F}(\mathbf{x}) \Leftrightarrow \begin{cases} \dot{x}_1 = F_1(x_1, x_2, ..., x_n) \\ \dot{x}_2 = F_2(x_1, x_2, ..., x_n) \\ \vdots \\ \dot{x}_n = F_n(x_1, x_2, ..., x_n) \end{cases} \quad \textbf{(S1)}$$

for $t \geq 0$ and with the initial conditions $\mathbf{x}(0) = \mathbf{x}_0$. Here, $\dot{x}_k = \frac{dx_k}{dt}$ and $\mathbf{F}(\mathbf{x})$ is a nonlinear vector function of $\mathbf{x}$. Then, using a first-order Taylor's expansion around a homogeneous steady state $\mathbf{x}^*$, we may rewrite $\mathbf{F}(\mathbf{x})$ as

$$\mathbf{F}(\mathbf{x}) \simeq \underbrace{\mathbf{F}(\mathbf{x}^*)}_{=0} + \mathbf{J}(\mathbf{x}^*)(\mathbf{x} - \mathbf{x}^*), \quad \textbf{(S2)}$$

where $\mathbf{J}(\mathbf{x}^*)$ is the Jacobian matrix of $\mathbf{F}$, given by

$$\mathbf{J}(\mathbf{x}) = \begin{pmatrix} \frac{\partial F_1}{\partial x_1} & \frac{\partial F_1}{\partial x_2} & \cdots & \frac{\partial F_1}{\partial x_n} \\ \frac{\partial F_2}{\partial x_1} & \frac{\partial F_2}{\partial x_2} & \cdots & \frac{\partial F_2}{\partial x_n} \\ \vdots & \vdots & \ddots & \vdots \\ \frac{\partial F_n}{\partial x_1} & \frac{\partial F_n}{\partial x_2} & \cdots & \frac{\partial F_n}{\partial x_n} \end{pmatrix}. \quad \textbf{(S3)}$$

We now want to deduce a linear approximation to fixed points. By setting $\tilde{\mathbf{x}} = \mathbf{x} - \mathbf{x}^*$, Eq. (S1) becomes

$$\frac{d\tilde{\mathbf{x}}}{dt} \simeq \mathbf{J}(\mathbf{x}^*)(\mathbf{x} - \mathbf{x}^*) = \mathbf{J}(\mathbf{x}^*)\tilde{\mathbf{x}} \quad \textbf{(S4)}$$

for $t \geq 0$, with initial conditions $\tilde{\mathbf{x}}(0) = \mathbf{x}_0 - \mathbf{x}^*$, and $\mathbf{J}(\mathbf{x}^*)$ is a constant matrix independent of $\mathbf{x}$. Hence, motivated by the one-dimensional linear case, the general solution is given by

$$\mathbf{x}(t) = \mathbf{x}^* + \sum_{i=1}^{n} a_k \mathbf{v}_k e^{\lambda_k t}, \quad \textbf{(S5)}$$

where $a_k$ are constants determined by the initial conditions, and $\lambda_k$ and $\mathbf{v}_k$ are the eigenvalues and eigenvectors of $\mathbf{J}(\mathbf{x}^*)$, respectively. The analysis is then based on the following steps

1. Determine the fixed point vector, $\mathbf{x}^*$, solving $\mathbf{F}(\mathbf{x}^*) = 0$;
2. Construct the Jacobian matrix $\mathbf{J}(\mathbf{x}^*)$;
3. Compute eigenvalues of $\mathbf{J}(\mathbf{x}^*)$ by solving the characteristic equation

$$\det|\mathbf{J}(\mathbf{x}^*) - \lambda \mathbf{I}| = 0, \quad \textbf{(S6)}$$

   where $\mathbf{I}$ is the $n \times n$ identity matrix.

Once these are achieved, stability or instability of a homogeneous steady state $\mathbf{x}^*$ depends on the real parts of the eigenvalues, Re$(\lambda)$, as follows

1. All eigenvalues have negative real parts $\Rightarrow$ $\mathbf{x}^*$ is locally stable;
2. At least one of the eigenvalues has a positive real part $\Rightarrow$ $\mathbf{x}^*$ is unstable;
3. Otherwise, there is no conclusion and it requires an investigation of the higher-order terms.

Studying the eigenvalues allows for stability analysis of linear dynamical systems and local stability analysis of nonlinear dynamical systems. In the following, we discuss applications of these linear stability methods in the case of two different species in various lattice configurations. The goal of this section is, again, to present the stepping stones behind the analytical results in the main text.

### Autonomous systems

In this section we study the simple case of a cell with two species $x$ and $y$, in which concentrations are governed by

$$\dot{x} = F_x(x,y) \tag{S7}$$
$$\dot{y} = F_y(x,y), \tag{S8}$$

where $F_x$ and $F_y$ are, for now, general nonlinear functions of two variables. The simplest kind of solutions for which linear stability can be studied are the constant solutions $(x^*,y^*)$, leading to the homogeneous system

$$0 = F_x(x^*,y^*) \tag{S9}$$
$$0 = F_y(x^*,y^*). \tag{S10}$$

Assuming we have identified a constant solution $(x^*,y^*)$, we can investigate whether this solution becomes unstable to tiny (small magnitude) perturbations by deriving and studying the linear constant-coefficient evolution equations for the perturbations of each constant concentration. Hence, we introduce two separate perturbations $\tilde{x}$ and $\tilde{y}$ for each constant concentration to get, as suggested in the previous section,

$$x(t) = x^* + \tilde{x}(t) \tag{S11}$$
$$y(t) = y^* + \tilde{y}(t). \tag{S12}$$

Substituting Eq. (S11)-Eq. (S12) into equations Eq. (S7)-Eq. (S8) leads to

$$\frac{d}{dt}\begin{pmatrix} x^* + \tilde{x} \\ y^* + \tilde{y} \end{pmatrix} = \frac{d}{dt}\begin{pmatrix} \tilde{x} \\ \tilde{y} \end{pmatrix} = \begin{pmatrix} F_x(x^* + \tilde{x}, y^* + \tilde{y}) \\ F_y(x^* + \tilde{x}, y^* + \tilde{y}) \end{pmatrix}. \tag{S13}$$

Since the perturbations $\tilde{x}$ and $\tilde{y}$ are assumed to be small, we may Taylor-expand the right sides of Eq. (S13) to low-order powers of $\tilde{x}$ and $\tilde{y}$. Hence, performing a first-order Taylor expansion of the dynamical system around $(x^*,y^*)$ leads to

$$\frac{d}{dt}\begin{pmatrix} \tilde{x} \\ \tilde{y} \end{pmatrix} \simeq \mathbf{J}(x^*,y^*)\begin{pmatrix} \tilde{x} \\ \tilde{y} \end{pmatrix}, \tag{S14}$$

where $\tilde{x} = x - x^*$, $\tilde{y} = y - y^*$ and $\mathbf{J} \equiv \mathbf{J}(x^*,y^*)$ is the Jacobian of $F = (F_x, F_y)$ evaluated at the steady state, given by

$$\mathbf{J} = \left.\begin{pmatrix} \frac{\partial F_x}{\partial x} & \frac{\partial F_x}{\partial y} \\ \frac{\partial F_y}{\partial x} & \frac{\partial F_y}{\partial y} \end{pmatrix}\right|_{(x^*,y^*)}. \tag{S15}$$

This yields a system of constant-coefficient linear differential equations, with the constant coefficients given by $\mathbf{J}$, which means that they depend on the choice of the constant solution $(x^*,y^*)$. Therefore we expect the general solution $(\tilde{x},\tilde{y})$ to be a superposition of particular solutions consisting of exponentials, given by

$$\begin{pmatrix} \tilde{x} \\ \tilde{y} \end{pmatrix} = C^+\mathbf{v}^+e^{\lambda^+t} + C^-\mathbf{v}^-e^{\lambda^-t}, \tag{S16}$$

where $\lambda^\pm$ and $\mathbf{v}^\pm$ are the eigenvalues and eigenvectors of the matrix $\mathbf{J}$. $C^+$ and $C^-$ are constants that depend on the initial conditions of the system (specifically the initial noise) and are obtained by solving the system

$$\begin{pmatrix} \tilde{x}(0) \\ \tilde{y}(0) \end{pmatrix} = C^+\mathbf{v}^+ + C^-\mathbf{v}^- \Leftrightarrow C^\pm = \frac{\tilde{x}(0)v_2^\mp - \tilde{y}(0)v_1^\mp}{v_1^\pm v_2^\mp - v_1^\mp v_2^\pm}, \tag{S17}$$

where $\mathbf{v}^\pm = (v_1^\pm, v_2^\pm)^T$. The real part of the eigenvalues $\lambda^\pm$ determines whether some perturbation grows or decreases in magnitude over time. We call the real parts of the eigenvalues the *growth rates*. If $\mathrm{Re}(\lambda^\pm) < 0$, then the homogeneous steady state $(x^*,y^*)$ is linearly stable. The growth rates can be obtained by finding the roots of the characteristic polynomial $p(\lambda)$, which in turn is obtained by setting the following determinant to zero

$$0 = p(\lambda) = \det(\mathbf{J} - \lambda\mathbf{I}) = (\mathbf{J}_{11} - \lambda)(\mathbf{J}_{22} - \lambda) - \mathbf{J}_{12}\mathbf{J}_{21} = \lambda^2 - \tau\lambda + \sigma, \tag{S18}$$

where we define

$$\tau = \mathrm{Tr}(\mathbf{J}) = \mathbf{J}_{11} + \mathbf{J}_{22} \tag{S19}$$
$$\sigma = \det(\mathbf{J}) = \mathbf{J}_{11}\mathbf{J}_{22} - \mathbf{J}_{12}\mathbf{J}_{21} \tag{S20}$$

and $\mathbf{J}_{ij}$ is the $(i,j)$ entry of matrix $\mathbf{J}$. A quadratic equation like Eq. (S18) has generally two distinct roots and so there are two eigenvalues $\lambda^+$ and $\lambda^-$ associated with the linear stability problem. These are given by

$$\lambda^{\pm} = \frac{\tau \pm \sqrt{\tau^2 - 4\sigma}}{2}. \tag{S21}$$

There are now two cases to consider regarding the discriminant $\tau^2 - 4\sigma$

- If $\tau^2 - 4\sigma < 0$, then $\lambda^{\pm}$ are complex numbers. Hence, we may write

$$\lambda^{\pm} = \frac{\tau}{2} \pm \mathrm{i}\frac{\sqrt{4\sigma - \tau^2}}{2}. \tag{S22}$$

  Since, from Eq. (S16), only the real part of a growth rate $\lambda$ can change the magnitude of $e^{\lambda t}$, and the two complex roots have the same real part $\tau/2$, we conclude that linear stability of the constant solution requires $\tau < 0$. Furthermore, $\sigma > \tau^2/4 > 0$.

- If $\tau^2 - 4\sigma \geq 0$, both eigenvalues are real and linear stability of the constant solution requires both roots (growth rates) to be negative numbers. The smaller root

$$\lambda^{-} = \frac{\tau - \sqrt{\tau^2 - 4\sigma}}{2} \tag{S23}$$

  will be negative if $\tau < 0$. Considering the larger root

$$\lambda^{+} = \frac{\tau + \sqrt{\tau^2 - 4\sigma}}{2} \tag{S24}$$

  we see that in order to be negative we need to have $\tau < 0$ and $|\tau| > \sqrt{\tau^2 - 4\sigma}$ which will hold if $\sigma > 0$.

Hence, we may say that a constant solution $(x^*, y^*)$ of the two coupled evolution equations Eq. (S7)-Eq. (S8) is linearly stable if and only if $\tau < 0$ and $\sigma > 0$. These are necessary and sufficient conditions for both eigenvalues $\lambda^+$ and $\lambda^-$ to have negative real parts. If either of these two conditions is violated, the constant solution becomes linearly unstable and will evolve into some new solution.
In general, and as previously mentioned, an equilibrium point $\mathbf{x}^*$ of the differential system is stable if all the eigenvalues of $\mathbf{J}$ have negative real parts. On the other hand, the equilibrium point is unstable if at least one of the eigenvalues has a positive real part. Note that no conclusion can be made if some of the eigenvalues have zero real parts while the others are all negative. This case can not be decided based on LSA. The nonlinear terms we left out in the Taylor expansion will, in this case, determine stability.

#### One-dimensional array

We now extend the previous discussion by looking at the one-dimensional periodic array consisting of $N$ cells with two species $x$ and $y$, in which concentrations in cell $j$ are governed by

$$\dot{x}_j = F_x(x_j, \{x_r\}, y_j, \{y_r\}) \tag{S25}$$
$$\dot{y}_j = F_y(x_j, \{x_r\}, y_j, \{y_r\}), \tag{S26}$$

where $\{x_r\}$ and $\{y_r\}$ are the sets of all the variables corresponding to cells that are not cell $j$. That is, $r \neq j$. We use $j$ as the focal cell index instead of $i$ to avoid confusion with the imaginary number i, though indexation with $i$ is interchangeably used in the following sections. $F_x$ and $F_y$ are, again, general nonlinear functions. Eq. (S25)-Eq. (S26) is translationally invariant, implying that the system written in matrix form, for all cells, contains a banded circulant matrix. Note that we have a total of $2N$ equations. As before, a homogeneous steady state of the system, $(x^*, y^*)$, requires $\dot{x}_j = \dot{y}_j = 0$ when $(x_j, y_j) = (x^*, y^*)$, for any cell $j$. This results in the homogeneous system

$$0 = F_x(x^*, \{x^*\}, y^*, \{y^*\}) \tag{S27}$$
$$0 = F_y(x^*, \{x^*\}, y^*, \{y^*\}). \tag{S28}$$

Next, we perturb this solution by setting $(x_j, y_j) = (x^* + \tilde{x}_j, y^* + \tilde{y}_j)$ and, following linearisation, we get

$$\frac{d}{dt}\begin{pmatrix}\tilde{x}_j \\ \tilde{y}_j\end{pmatrix} \simeq \mathbf{J}\begin{pmatrix}\tilde{x}_j \\ \tilde{y}_j\end{pmatrix} + \sum_{r \neq j} \mathbf{J}^r \begin{pmatrix}\tilde{x}_r \\ \tilde{y}_r\end{pmatrix}, \tag{S29}$$

where

$$\mathbf{J} = \begin{pmatrix} \frac{\partial F_x}{\partial x_j} & \frac{\partial F_x}{\partial y_j} \\ \frac{\partial F_y}{\partial x_j} & \frac{\partial F_y}{\partial y_j} \end{pmatrix}\Bigg|_{(x^*,y^*)} \text{ and } \mathbf{J}^r = \begin{pmatrix} \frac{\partial F_x}{\partial x_r} & \frac{\partial F_x}{\partial y_r} \\ \frac{\partial F_y}{\partial x_r} & \frac{\partial F_y}{\partial y_r} \end{pmatrix}\Bigg|_{(x^*,y^*)}. \tag{S30}$$

Expanding the previous expressions, we rewrite the linearised system as

$$\frac{d}{dt}\begin{pmatrix} \tilde{x}_j \\ \tilde{y}_j \end{pmatrix} = \begin{pmatrix} \mathbf{J}_{11}\tilde{x}_j + \mathbf{J}_{12}\tilde{y}_j + \sum_{r\neq j}[\mathbf{J}^r_{11}\tilde{x}_r + \mathbf{J}^r_{12}\tilde{y}_r] \\ \mathbf{J}_{21}\tilde{x}_j + \mathbf{J}_{22}\tilde{y}_j + \sum_{r\neq j}[\mathbf{J}^r_{21}\tilde{x}_r + \mathbf{J}^r_{22}\tilde{y}_r] \end{pmatrix}. \tag{S31}$$

Next, we decouple the system of $2N$ equations by performing a discrete Fourier transform with respect to $j$ and changing the variables as follows, for $1 \le q \le N$,

$$\tilde{x}_j = \sum_{q=1}^{N} \xi_q e^{2\pi \mathrm{i} qj/N} \tag{S32}$$

$$\tilde{y}_j = \sum_{q=1}^{N} \eta_q e^{2\pi \mathrm{i} qj/N}, \tag{S33}$$

which may also be written as

$$\xi_q = \frac{1}{N}\sum_{j=1}^{N} \tilde{x}_j e^{-2\pi \mathrm{i} qj/N} \tag{S34}$$

$$\eta_q = \frac{1}{N}\sum_{j=1}^{N} \tilde{y}_j e^{-2\pi \mathrm{i} qj/N}. \tag{S35}$$

Eq. (S32) and Eq. (S33) are Fourier transforms in the *exponential form* (Bracewell and Bracewell, 1986). In order to rewrite the system for $(\xi_q, \eta_q)$, start by differentiating $\xi_q$ and substituting in Eq. (S31), as follows, with $\bar{q} \equiv q/N$,

$$\frac{d\xi_q}{dt} = \frac{1}{N}\sum_{j=1}^{N} \frac{d\tilde{x}_j}{dt} e^{-2\pi \mathrm{i}\bar{q}j} \tag{S36}$$

$$= \frac{1}{N}\sum_{j=1}^{N}\left[\mathbf{J}_{11}\tilde{x}_j + \mathbf{J}_{12}\tilde{y}_j + \sum_{r\neq j}[\mathbf{J}^r_{11}\tilde{x}_r + \mathbf{J}^r_{12}\tilde{y}_r]\right] e^{-2\pi \mathrm{i}\bar{q}j} \tag{S37}$$

$$= \frac{\mathbf{J}_{11}}{N}\sum_{j=1}^{N}\tilde{x}_j e^{-2\pi \mathrm{i}\bar{q}j} + \frac{\mathbf{J}_{12}}{N}\sum_{j=1}^{N}\tilde{y}_j e^{-2\pi \mathrm{i}\bar{q}j} + \frac{1}{N}\sum_{j=1}^{N}\sum_{r\neq j}[\mathbf{J}^r_{11}\tilde{x}_r + \mathbf{J}^r_{12}\tilde{y}_r]\, e^{-2\pi \mathrm{i}\bar{q}j} \tag{S38}$$

$$= \mathbf{J}_{11}\xi_q + \mathbf{J}_{12}\eta_q + \frac{1}{N}\sum_{j=1}^{N}\sum_{r\neq j}[\mathbf{J}^r_{11}\tilde{x}_r + \mathbf{J}^r_{12}\tilde{y}_r]\, e^{-2\pi \mathrm{i}\bar{q}j}, \tag{S39}$$

where we used the definitions in Eq. (S34) and Eq. (S35). The third term of Eq. (S39) requires more thought. The goal is to write it in terms of $\xi_q$ exclusively. Let us focus on the terms involving $\tilde{x}_r$ first. Take $r = j + \Delta j$, $\Delta j \in S$, where $S$ is the set of all integers such that $j + \Delta j$ is an index of a cell and $\Delta j \neq 0$. Notice that the indexation of $\mathbf{J}^r_{11} = \mathbf{J}^{j+\Delta j}_{11}$ is always relative to $\Delta j$ and therefore constant on $j$ ($F_x$ and $F_y$ do not change with $j$). Thus we may take $\mathbf{J}^r_{11} \equiv \mathbf{J}^{\Delta j}_{11}$ and write

$$\frac{1}{N}\sum_{j=1}^{N}\sum_{r\neq j}\mathbf{J}^r_{11}\tilde{x}_r e^{-2\pi \mathrm{i}\bar{q}j} = \sum_{\Delta j\in S}\mathbf{J}^{\Delta j}_{11}\frac{1}{N}\sum_{j=1}^{N}\tilde{x}_{j+\Delta j}e^{-2\pi \mathrm{i}\bar{q}j} = \sum_{\Delta j\in S}\mathbf{J}^{\Delta j}_{11}\xi_q e^{-2\pi \mathrm{i}\bar{q}\Delta j}. \tag{S40}$$

The last equality is obtained by seeing that, excluding the sum over $\Delta j$, we have

$$\frac{1}{N}\sum_{j=1}^{N}\tilde{x}_{j+\Delta j}e^{-2\pi \mathrm{i}\bar{q}j} = \frac{e^{-2\pi \mathrm{i}\bar{q}\Delta j}}{N}\sum_{j=1}^{N}\tilde{x}_{j+\Delta j}e^{-2\pi \mathrm{i} q(j+\Delta j)/N} \tag{S41}$$

$$= \frac{e^{-2\pi \mathrm{i}\bar{q}\Delta j}}{N}\sum_{j=1}^{N}\tilde{x}_j e^{-2\pi \mathrm{i}\bar{q}j} \tag{S42}$$

$$= \xi_q e^{-2\pi i \bar{q}\Delta j}, \tag{S43}$$

where we used the fact that we are working on the $\mathbb{Z}/N$ ring, thus $\tilde{x}_j = \tilde{x}_{j+N}$ and the sum term is $N$-periodic. We apply a similar analysis to $\tilde{y}$. This shows that any term involving $\tilde{x}_{j+\Delta j}$ or $\tilde{y}_{j+\Delta j}$, $\forall \Delta j \in S$, can be transformed into a linear term involving $\xi_q$ or $\eta_q$, respectively. Hence, we have that

$$\frac{1}{N}\sum_{j=1}^{N}\sum_{r\neq j}[\mathbf{J}_{11}^{r}\tilde{x}_r + \mathbf{J}_{12}^{r}\tilde{y}_r]\, e^{-2\pi i\bar{q}j} = \sum_{\Delta j\in S}\left[\mathbf{J}_{11}^{\Delta j}\xi_q + \mathbf{J}_{12}^{\Delta j}\eta_q\right] e^{-2\pi i\bar{q}\Delta j} \tag{S44}$$

$$= \xi_q \sum_{\Delta j\in S}\mathbf{J}_{11}^{\Delta j} e^{-2\pi i\bar{q}\Delta j} + \eta_q \sum_{\Delta j\in S}\mathbf{J}_{12}^{\Delta j} e^{-2\pi i\bar{q}\Delta j}. \tag{S45}$$

We may say that such linearisation leads to four functions that take into account the spatial coupling terms of each equation, depending solely on $q$. In matrix form, such functions can be captured by

$$\mathbf{\Omega}_{\bar{q}} \equiv \mathbf{\Omega}_{\bar{q}}(q) = \sum_{\Delta j\in S}\begin{pmatrix}\mathbf{J}_{11}^{\Delta j}e^{-2\pi i\bar{q}\Delta j} & \mathbf{J}_{12}^{\Delta j}e^{-2\pi i\bar{q}\Delta j}\\ \mathbf{J}_{21}^{\Delta j}e^{-2\pi i\bar{q}\Delta j} & \mathbf{J}_{22}^{\Delta j}e^{-2\pi i\bar{q}\Delta j}\end{pmatrix} = \sum_{\Delta j\in S}\mathbf{J}^{\Delta j}e^{-2\pi i\bar{q}\Delta j}. \tag{S46}$$

In general, it is then possible to rewrite Eq. (S25)-Eq. (S26) with respect to $(\xi_q, \eta_q)$ to get the following system of constant-coefficient linear differential equations

$$\dot{\xi}_q = F_\xi(\xi_q,\eta_q) \simeq (\mathbf{J}_{11} + \mathbf{\Omega}_{\bar{q}11})\xi_q + (\mathbf{J}_{12} + \mathbf{\Omega}_{\bar{q}12})\eta_q \tag{S47}$$

$$\dot{\eta}_q = F_\eta(\xi_q,\eta_q) \simeq (\mathbf{J}_{21} + \mathbf{\Omega}_{\bar{q}21})\xi_q + (\mathbf{J}_{22} + \mathbf{\Omega}_{\bar{q}22})\eta_q \tag{S48}$$

or simply

$$\frac{d}{dt}\begin{pmatrix}\xi_q\\ \eta_q\end{pmatrix} \simeq \mathbf{L}_{\bar{q}}\begin{pmatrix}\xi_q\\ \eta_q\end{pmatrix}, \tag{S49}$$

where $\mathbf{L}_{\bar{q}} = \mathbf{J} + \mathbf{\Omega}_{\bar{q}}$. Following the previous section, the general solution of Eq. (S49) is then given by

$$\begin{pmatrix}\xi_q\\ \eta_q\end{pmatrix} = C_{\bar{q}}^{+}\mathbf{v}_{\bar{q}}^{+}e^{\lambda_{\bar{q}}^{+}t} + C_{\bar{q}}^{-}\mathbf{v}_{\bar{q}}^{-}e^{\lambda_{\bar{q}}^{-}t}, \tag{S50}$$

where $\lambda_{\bar{q}}^{\pm}$ and $\mathbf{v}_{\bar{q}}^{\pm}$ are the eigenvalues and eigenvectors of the matrix $\mathbf{L}_{\bar{q}}$. $C_{\bar{q}}^{+}$ and $C_{\bar{q}}^{-}$ are constants depending on the initial conditions of the problem (note that they are different for each $q$, since each yields a separate system). We have

$$\lambda_{\bar{q}}^{\pm} = \frac{\tau_{\bar{q}} \pm \sqrt{\tau_{\bar{q}}^2 - 4\sigma_{\bar{q}}}}{2} \tag{S51}$$

$$= \frac{1}{2}\left(\mathbf{L}_{\bar{q}11} + \mathbf{L}_{\bar{q}22} \pm \sqrt{(\mathbf{L}_{\bar{q}11} + \mathbf{L}_{\bar{q}22})^2 - 4(\mathbf{L}_{\bar{q}11}\mathbf{L}_{\bar{q}22} - \mathbf{L}_{\bar{q}12}\mathbf{L}_{\bar{q}21})}\right), \tag{S52}$$

where, in this case,

$$\tau_{\bar{q}} = \mathrm{Tr}(\mathbf{L}_{\bar{q}}) = \mathbf{L}_{\bar{q}11} + \mathbf{L}_{\bar{q}22} \tag{S53}$$

$$\sigma_{\bar{q}} = \det(\mathbf{L}_{\bar{q}}) = \mathbf{L}_{\bar{q}11}\mathbf{L}_{\bar{q}22} - \mathbf{L}_{\bar{q}12}\mathbf{L}_{\bar{q}21}. \tag{S54}$$

The solution of Eq. (S29) follows by applying the variable change Eq. (S32)-Eq. (S33) to the previous solution, leading to

$$\begin{pmatrix}\tilde{x}_j\\ \tilde{y}_j\end{pmatrix} = \sum_{q=1}^{N}\left[C_{\bar{q}}^{+}\mathbf{v}_{\bar{q}}^{+}e^{\lambda_{\bar{q}}^{+}t} + C_{\bar{q}}^{-}\mathbf{v}_{\bar{q}}^{-}e^{\lambda_{\bar{q}}^{-}t}\right]e^{2\pi i\bar{q}j}. \tag{S55}$$

Again, we argue that a homogeneous steady state $(x^*, y^*)$ is linearly stable if and only if Re$(\lambda_{\bar{q}}^{\pm}) < 0$, $\forall q$. Further details on conditions for stability were discussed in the previous section. We are now interested in studying what happens to the patterning solutions as $t \to \infty$. The real parts of the eigenvalues characterise the exponential growth rate along the eigenvectors, and so, if Re$(\lambda_{\bar{q}}^{\pm}) < 0$, $\forall \bar{q}$, perturbations do not grow and the homogeneous solution is linearly stable. In contrast, the maximum value of Re$(\lambda_{\bar{q}}^{\pm})$ indicates the fastest growing mode or wavelength $\bar{q}$ at the onset of instability. When such a value is positive, the homogeneous state is linearly unstable, and a pattern with the characteristic wavelength of the set of modes $\bar{q}$ that maximise Re$(\lambda^{\pm}_{\bar{q}})$ arises for small $t$. For large $t$, we expect a dominant pattern to emerge and a consequent simplification of Eq. (S55) given by the superposition of the fastest growing modes, although this may not be true if nonlinearities determine emergent patterning. This translates into a maximisation problem over all $\bar{q}$ (note that $N$, in this case, is simply a refinement of the discretisation and, therefore, does not affect the maximisation problem). That is, we want to solve

$$\max_{\bar{q}} \text{Re}(\lambda^{+}_{\bar{q}}) = \max_{\bar{q}} \text{Re}\left[\frac{\tau_{\bar{q}} + \sqrt{\tau_{\bar{q}}^2 - 4\sigma_{\bar{q}}}}{2}\right] \tag{S56}$$

$$= \frac{1}{2}\max_{\bar{q}} \text{Re}\left[\mathbf{L}_{\bar{q}11} + \mathbf{L}_{\bar{q}22} + \sqrt{(\mathbf{L}_{\bar{q}11} + \mathbf{L}_{\bar{q}22})^2 - 4(\mathbf{L}_{\bar{q}11}\mathbf{L}_{\bar{q}22} - \mathbf{L}_{\bar{q}12}\mathbf{L}_{\bar{q}21})}\right]. \tag{S57}$$

Since $\mathbf{J}$ is independent of $\bar{q}$, we only need to look into optimising the terms involving $\mathbf{\Omega}_{\bar{q}}$. This may be hard in general, but, despite the complexity in Eq. (S57), the coupling terms regard only the terms $\mathbf{\Omega}_{\bar{q}12}$ and $\mathbf{\Omega}_{\bar{q}21}$ in many applications, as we will see in the example in Example 1.1 and the case of Notch-Delta below. Once the set of fastest-growing modes (maximisers) $\mathcal{W}$ is obtained, the sum in Eq. (S55) is restricted to this set and becomes, asymptotically,

$$\begin{pmatrix}\tilde{x}_j \\ \tilde{y}_j\end{pmatrix} \approx \mathbf{v}^* e^{\lambda^* t} \sum_{\bar{q}\in\mathcal{W}} C^{+}_{\bar{q}} e^{2\pi \mathrm{i} \bar{q} j}, \tag{S58}$$

where $\lambda^*$ is the growth rate corresponding to the fastest growing mode (eigenvalue with the largest real part) and $\mathbf{v}^*$ its corresponding eigenvector. Small perturbations grow exponentially (without temporal oscillations) on a time scale of order $1/\text{Re}(\lambda^*)$.

**Two-dimensional array**

Consider now a $N \times M$ two-dimensional lattice and assume each cell is labelled by $(j,k)$, with $1 \le j \le N, 1 \le k \le M$. We ignore the lattice shape for now. Again, we study the dynamics of two species $x$ and $y$. Notice first that the main difference between this case and the one-dimensional array studied before is that now our decoupling and change of variables must account for two indices as well. Therefore, we relabel $j \equiv (j,k)$ in our previous analysis and aim to solve the system

$$\dot{x}_{j,k} = F_x(x_{j,k}, \{x_r\}, y_{j,k}, \{y_r\}) \tag{S59}$$

$$\dot{y}_{j,k} = F_y(x_{j,k}, \{x_r\}, y_{j,k}, \{y_r\}), \tag{S60}$$

where $r \neq (j,k)$. Linearisation leads to

$$\frac{d}{dt}\begin{pmatrix}\tilde{x}_{j,k} \\ \tilde{y}_{j,k}\end{pmatrix} \simeq \mathbf{J}\begin{pmatrix}\tilde{x}_{j,k} \\ \tilde{y}_{j,k}\end{pmatrix} + \sum_{r\neq(j,k)} \mathbf{J}^r \begin{pmatrix}\tilde{x}_r \\ \tilde{y}_r\end{pmatrix}. \tag{S61}$$

Using the two-dimensional Fourier transform, we may change the variables to get, for $1 \le q \le N$ and $1 \le p \le M$,

$$\tilde{x}_{j,k} = \sum_{q=1}^{N}\sum_{p=1}^{M} \xi_{q,p} e^{2\pi \mathrm{i}(qj/N + pk/M)} \tag{S62}$$

$$\tilde{y}_{j,k} = \sum_{q=1}^{N}\sum_{p=1}^{M} \eta_{q,p} e^{2\pi \mathrm{i}(qj/N + pk/M)}, \tag{S63}$$

with inverted transform given by

$$\xi_{q,p} = \frac{1}{MN}\sum_{k=1}^{M}\sum_{j=1}^{N} \tilde{x}_{j,k} e^{-2\pi \mathrm{i}(qj/N + pk/M)} \tag{S64}$$

$$\eta_{q,p} = \frac{1}{MN}\sum_{k=1}^{M}\sum_{j=1}^{N} \tilde{y}_{j,k} e^{-2\pi \mathrm{i}(qj/N + pk/M)}. \tag{S65}$$

Following the steps in the previous section, we easily see that the spatial coupling function is now given by

$$\mathbf{\Omega}_{\bar{q},\bar{p}} \equiv \mathbf{\Omega}_{\bar{q},\bar{p}}(q,p) = \sum_{\Delta jk\in S} \mathbf{J}^{\Delta jk} e^{-2\pi \mathrm{i}(\bar{q}\Delta j + \bar{p}\Delta k)}, \tag{S66}$$

where $(\bar{q},\bar{p}) = (q/N, p/M)$ and $\Delta jk = (\Delta j, \Delta k)$. In this case, we took $r = (j+\Delta j, k+\Delta k)$, $(\Delta j, \Delta k) \in S$, where $S$ is now the set of all pairs of integers such that $(j+\Delta j, k+\Delta k)$ is an index of a cell and $(\Delta j, \Delta k) \neq (0,0)$. $\mathbf{J}^{\Delta jk}$ is given, as before, by Eq. (S30). One interesting aspect of this generalised analysis is that it is independent of the lattice shape, as long as the connectivity matrix corresponds to a regular graph ($S$ is independent of indexes). Notice that a generalisation to any spatial dimension follows easily from this step. The particular case when weighting is symmetric is discussed in Example 1.1.
We then obtain the system

$$\frac{d}{dt}\begin{pmatrix}\xi_{q,p}\\ \eta_{q,p}\end{pmatrix} \simeq \mathbf{L}_{\bar{q},\bar{p}}\begin{pmatrix}\xi_{q,p}\\ \eta_{q,p}\end{pmatrix}, \tag{S67}$$

where $\mathbf{L}_{\bar{q},\bar{p}} = \mathbf{J} + \mathbf{\Omega}_{\bar{q},\bar{p}}$. The general solution of Eq. (S67) is then given by

$$\begin{pmatrix}\xi_{q,p}\\ \eta_{q,p}\end{pmatrix} = C^{+}_{\bar{q},\bar{p}} e^{\lambda^{+}_{\bar{q},\bar{p}} t}\mathbf{v}^{+}_{\bar{q},\bar{p}} + C^{-}_{\bar{q},\bar{p}} e^{\lambda^{-}_{\bar{q},\bar{p}} t}\mathbf{v}^{-}_{\bar{q},\bar{p}}, \tag{S68}$$

where $\lambda^{\pm}_{\bar{q},\bar{p}}$ and $\mathbf{v}^{\pm}_{\bar{q},\bar{p}}$ are the eigenvalues and eigenvectors of the matrix $\mathbf{L}_{\bar{q},\bar{p}}$. $C^{+}_{\bar{q},\bar{p}}$ and $C^{-}_{\bar{q},\bar{p}}$ are constants depending on the initial conditions of the problem. The solution to Eq. (S61) is then given by

$$\begin{pmatrix}\tilde{x}_{j,k}\\ \tilde{y}_{j,k}\end{pmatrix} = \sum_{q=1}^{N}\sum_{p=1}^{M}\left[C^{+}_{\bar{q},\bar{p}} e^{\lambda^{+}_{\bar{q},\bar{p}} t}\mathbf{v}^{+}_{\bar{q},\bar{p}} + C^{-}_{\bar{q},\bar{p}} e^{\lambda^{-}_{\bar{q},\bar{p}} t}\mathbf{v}^{-}_{\bar{q},\bar{p}}\right] e^{2\pi \mathrm{i}(\bar{q}j+\bar{p}k)}. \tag{S69}$$

Similar to the one-dimensional case, the fastest growing modes are obtained by maximizing the real part of the growth rates over $\bar{q}$ and $\bar{p}$, and the asymptotic behaviour of the solution of the linearised problem is dominated by the terms corresponding to the fastest growing modes

$$\begin{pmatrix}\tilde{x}_{j,k}\\ \tilde{y}_{j,k}\end{pmatrix} \approx \mathbf{v}^{*} e^{\lambda^{*} t}\sum_{(\bar{q},\bar{p})\in\mathcal{W}} C^{+}_{\bar{q},\bar{p}} e^{2\pi \mathrm{i}(\bar{q}j+\bar{p}k)}, \tag{S70}$$

where $\lambda^{*}$ is the growth rate corresponding to the fastest growing modes, which comprise $\mathcal{W}$, and $\mathbf{v}^{*}$ its corresponding eigenvector. When nonlinearities affect the solution, we do not expect, in general, solutions of the form Eq. (S70) to arise in the full nonlinear system. The two-dimensional system will be the main focus in this paper.

*Remark 1.1* (Continuum limit and reaction-diffusion systems) Feedback interactions between morphogens (understood as long-range diffusible ligands in many contexts (Lawrence and Struhl, 1996; Vincent and Briscoe, 2001; Tabata and Takei, 2004)) have previously been discussed by Alan Turing regarding skin patterning in his famous paper (Turing, 1952). Many of the patterns discussed in this work share similar features with the type of patterns expected from long-range signalling via filopodia (Meinhardt, 2003, 2008; Kondo, 2009). While diffusion is a linear process where the flux is proportional to the concentration gradient of morphogens, discrete lattice-based long-range signalling imposes nonlinear effects on patterning, often due to the production functions (Hill functions in our case). Nonetheless, as suggested in Binshtok and Sprinzak (2018); Hamada et al. (2014), the mathematical equivalence between the two approaches is worth noticing.
Consider the one-dimensional translationally invariant system given by

$$\frac{du_i}{dt} = f(u_i, v_{i-1}, v_{i+1}) \tag{S71}$$

$$\frac{dv_i}{dt} = g(u_i, v_i). \tag{S72}$$

By considering the continuum limit $u_i \to u(i\Delta)$ (and similarly for $v_i$), we get

$$\frac{\partial u(t,x)}{\partial t} = f(u(t,x), v(t,x-\Delta), v(t,x+\Delta)) \tag{S73}$$

$$\frac{\partial v(t,x)}{\partial t} = g(u(t,x), v(t,x)). \tag{S74}$$

Expanding to second order in $\Delta$ leads to

$$\frac{\partial u(t,x)}{\partial t} \simeq f(u(t,x), v(t,x) - \Delta\partial_x v + \frac{1}{2}\Delta^2\partial_{xx}v, v(t,x) + \Delta\partial_x v + \frac{1}{2}\Delta^2\partial_{xx}v) \tag{S75}$$

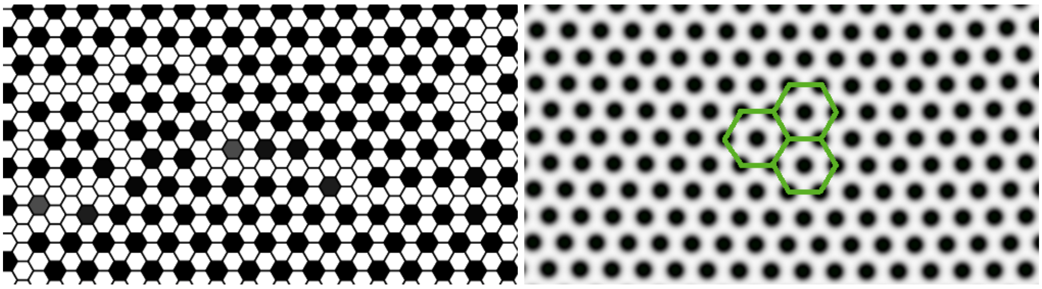

**(a)** **(b)**

**Figure S1. Reaction-diffusion systems mimic juxtacrine signalling.**
**(a)** Lateral inhibition patterning. **(b)** Soliton-type patterning via the Gray-Scott Model of a reaction-diffusion system resembles lateral inhibition on a hexagonal lattice (sketched in green). Here $F = 0.03$, $k = 0.062$ and $D_n = 2D_d$.

$$\frac{\partial v(t,x)}{\partial t} = g(u(t,x), v(t,x)). \tag{S76}$$

A consistent expansion of $f$ will lead to the reaction-diffusion form, possibly with a convection ($\partial_x$) term (in the general case). If, for example, $f(u_i, v_{i-1}, v_{i+1}) = v_{i-1} + v_{i+1}$, we may identify $v_{i-1} + v_{i+1} \sim 2v + h\partial_{xx}v$, for some small $h \equiv \Delta^2$.
Figure S1 compares a simulation of the Collier model with the Gray-Scott model of reaction-diffusion (Doelman et al., 1997; McGough and Riley, 2004), given by

$$\frac{\partial n}{\partial t} = -nd^2 + \mathcal{F}(1-n) + D_n \nabla n \tag{S77}$$

$$\frac{\partial d}{\partial t} = nd^2 - (\mathcal{F} + \kappa)d + D_d \nabla d, \tag{S78}$$

where $\mathcal{F}$ is the feed rate, $\kappa$ is the kill rate and $D_n$ and $D_d$ are the diffusion rates. Lateral inhibition is mimicked for specific ranges of such rates. Sparser patterns might be expected under similar regimes.

### Examples

So far, we have been discussing linear stability methods in a fairly general way. This section discusses coupling simplifications based on symmetry and presents an application to a one-dimensional system. The two-dimensional case will be studied in the particular case of Notch-Delta signalling.

*Example 1.1* (Simplification of coupling functions) In many cases, the coupling functions $\mathbf{\Omega}_{\bar{q}}$ can be simplified under specific conditions. In the one-dimensional ring, for example, if the dependence on $x_r$, $r \neq j$, is given as a sum of indexed terms corresponding to immediate neighbours, that is, $r \in \{j-1, j+1\}$ ($\Delta j = \pm 1$) and

$$F_x(x_j, \{x_r\}, y_j, \{y_r\}) \equiv F_x(x_{j-1} + x_{j+1}) \tag{S79}$$

$$F_y(x_j, \{x_r\}, y_j, \{y_r\}) \equiv F_y(x_j, y_j), \tag{S80}$$

then

$$\mathbf{J}^{\pm 1} = \begin{pmatrix} 1 & 0 \\ 0 & 0 \end{pmatrix} \tag{S81}$$

and thus the only non-zero term in $\mathbf{\Omega}_{\bar{q}}$ is

$$\mathbf{\Omega}_{\bar{q}11} = e^{2\pi i \bar{q}} + e^{-2\pi i \bar{q}} = 2\cos(2\pi\bar{q}). \tag{S82}$$

Generalising to the $m$th neighbour of cell $j$, that is, $\Delta j \in S^m \equiv \{\pm m\}$, it is not hard to see that,

$$\mathbf{\Omega}_{\bar{q}11} = 2\cos(2\pi\bar{q}m). \tag{S83}$$

Notice now that if we consider $F_x$ of the form $F_x\left(\sum_{\Delta j \in \Sigma S^m} x_{j+\Delta j}\right)$, with $\Sigma S^m = \{\pm 1, ..., \pm m\}$, then the coupling function is easily obtained by summing, up to $m$, the previous terms. Other simplifications are possible depending on the nature of $\mathbf{J}$.
In two dimensions, we have more interesting cases to look at. First, depending on the lattice shape, $(\Delta j, \Delta k)$ might mean different things regarding which cells are neighbours of a focal cell $(j,k)$. The analysis, however,

is independent of this. Thus we merely refer to lattice shapes as motivation for the choices of $S$, as mentioned before. In a squared lattice, one of two cases is often relevant: either a cell is affected by only 4 neighbours or all 8 (Figure S2a). In the first case, we take $S = \{(0,\pm 1),(\pm 1,0)\}$. Here, we use the notation $\Omega_{\bar{q},\bar{p}} \equiv [\boldsymbol{\Omega}_{\bar{q},\bar{p}}]_{11}$ as a simplification (it is irrelevant which variable is affected). Again, we take equally weighted sums of the neighbours' terms. In this case, we get, from Eq. (S66),

$$\Omega_{\bar{q},\bar{p}} = \sum_{\Delta j \in S} e^{-2\pi \mathrm{i}(\bar{q}\Delta j + \bar{p}\Delta k)} = 2\left[\cos(2\pi\bar{q}) + \cos(2\pi\bar{p})\right]. \tag{S84}$$

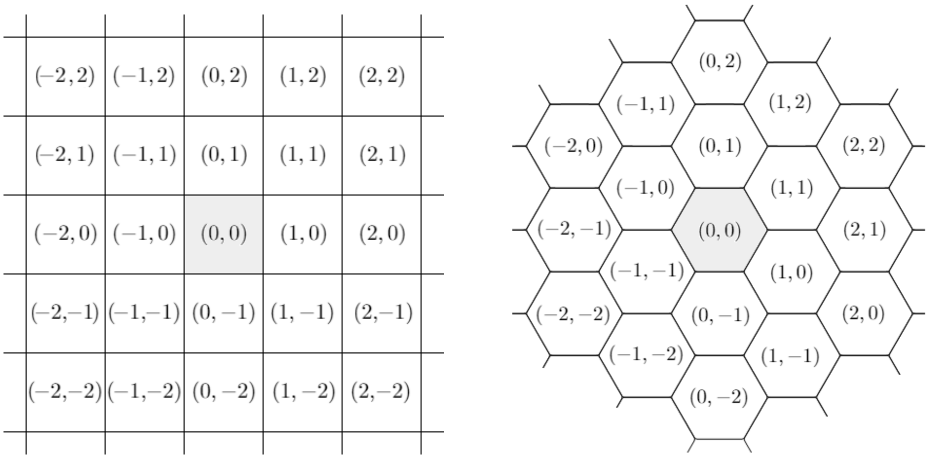


**(a)** Squared lattice

**(b)** Hexagonal lattice

**Figure S2. Cell lattices and relative indexation.**

In the case of the regular hexagonal lattice, we have several labelling possibilities. Using the labelling in Figure S2b, we get, for immediate neighbours, $S = \{(\pm 1,0),(0,\pm 1),(\pm 1,\pm 1)\}$ and

$$\Omega_{\bar{q},\bar{p}} = 2\left[\cos\left(2\pi\bar{q}\right) + \cos\left(2\pi\bar{p}\right) + \cos\left(2\pi\left(\bar{q}+\bar{p}\right)\right)\right]. \tag{S85}$$

More generally, if the coupling weights, captured by $\mathbf{J}^{\Delta jk}$, are symmetric, that is, $\mathbf{J}^{\Delta jk} = \mathbf{J}^{\Delta(-j)(-k)}$, then

$$\boldsymbol{\Omega}_{\bar{q},\bar{p}} = \sum_{\Delta jk \in S} \mathbf{J}^{\Delta jk} \cos(2\pi(\bar{q}\Delta j + \bar{p}\Delta k)). \tag{S86}$$

*Example 1.2* (Ring of cells) Consider two species $x$ and $y$ governed by the following equations on a ring of $N$ cells

$$\frac{dx_j}{dt} = f\left(\frac{y_{j-1}+y_{j+1}}{2}\right) + \mu x_j \tag{S87}$$

$$\frac{dy_j}{dt} = g(x_j) + \rho y_j, \tag{S88}$$

where $\mu, \rho \in \mathbb{R}$. Define

$$F_x(x_j, y_j, y_{j-1}, y_{j+1}) = f\left(\frac{y_{j-1}+y_{j+1}}{2}\right) + \mu x_j \tag{S89}$$

$$F_y(x_j, y_j, y_{j-1}, y_{j+1}) = g(x_j) + \rho y_j. \tag{S90}$$

Following linearisation around a homogeneous steady state, the relevant matrices are given by

$$\mathbf{J} = \begin{pmatrix} \mu & 0 \\ g'(x_j) & \rho \end{pmatrix}\Bigg|_{(x^*,y^*)} = \begin{pmatrix} \mu & 0 \\ g'(x^*) & \rho \end{pmatrix} \tag{S91}$$

$$\mathbf{J}^{\pm 1} = \begin{pmatrix} 0 & \frac{1}{2} f'\left(\frac{y_{j-1}+y_{j+1}}{2}\right) \\ 0 & 0 \end{pmatrix}\Bigg|_{(x^*,y^*)} = \begin{pmatrix} 0 & \frac{1}{2} f'(y^*) \\ 0 & 0 \end{pmatrix}. \tag{S92}$$

In this case, we have $S = \{-1, 1\}$ and $\mathbf{J}^1 = \mathbf{J}^{-1}$ (not to be confused with the inverse of a matrix). Thus we may use Eq. (S86) from Example 1.1 and write

$$\boldsymbol{\Omega}_{\bar{q}} = \sum_{\Delta j \in S} \mathbf{J}^{\Delta j} \cos(2\pi\bar{q}\Delta j) \tag{S93}$$

$$= \begin{pmatrix} 0 & f'(y^*)(\cos(2\pi\bar{q}) + \cos(-2\pi\bar{q})) \\ 0 & 0 \end{pmatrix} \tag{S94}$$

$$= \begin{pmatrix} 0 & 2f'(y^*)\cos(2\pi\bar{q}) \\ 0 & 0 \end{pmatrix} \tag{S95}$$

and so

$$\mathbf{L}_{\bar{q}} = \mathbf{J} + \mathbf{\Omega}_{\bar{q}} = \begin{pmatrix} \mu & 2f'(y^*)\cos(2\pi\bar{q}) \\ g'(x^*) & \rho \end{pmatrix}. \tag{S96}$$

We define the coupling term by $\Omega_{\bar{q}} \equiv 2\cos(2\pi\bar{q})$ (not to be confused with the matrix counterpart, $\mathbf{\Omega}_{\bar{q}}$, in bold) and let $A = f'(y^*)$ and $B = g'(x^*)$. In fact, in this case, $\Omega_{\bar{q}} = \mathbf{\Omega}_{\bar{q}12}/A$. The eigenvalues of $\mathbf{L}$ are then given by

$$\lambda_{\bar{q}}^{\pm} = \frac{1}{2}\left(\mathbf{L}_{\bar{q}11} + \mathbf{L}_{\bar{q}22} \pm \sqrt{(\mathbf{L}_{\bar{q}11} + \mathbf{L}_{\bar{q}22})^2 - 4(\mathbf{L}_{\bar{q}11}\mathbf{L}_{\bar{q}22} - \mathbf{L}_{\bar{q}12}\mathbf{L}_{\bar{q}21})}\right) \tag{S97}$$

$$= \frac{1}{2}\left(\mu + \rho \pm \sqrt{(\mu+\rho)^2 - 4(\mu\rho - AB\Omega_{\bar{q}})}\right). \tag{S98}$$

From the previous analysis, the homogeneous solution is linearly stable if and only if $\mu + \rho < 0$ and $\mu\rho > AB\Omega_{\bar{q}}$ for all $\bar{q}$. Maximising $\mathrm{Re}(\lambda_{\bar{q}}^{\pm})$ with respect to $\bar{q}$ relies then on maximising $AB\Omega_{\bar{q}}$, since $AB\Omega_{\bar{q}} > \mu\rho$ leads to a heterogeneous solution.

*Remark 1.2* (Euclidean and hexagonal distances)

For a focal cell $i$ ($R_0$), the recursive definition of the signalling rings $R_k$ as the set of cells that are immediate neighbours of cells in $R_{k-1}$ and are not in $\bigcup_{n=1}^{k-1} R_n$ can be a bit ambiguous regarding the realistic reach of protrusions and the regular hexagonal lattice. First, notice that a hexagonal-Manhattan-type distance, $d_H(i,j)$, provides an alternative definition of such sets in the following way: a cell $i$ is in $R_k$ if it is not in $\bigcup_{n=1}^{k-1} R_n$ and $d_H(i,j) = k$ (between cell centres and assuming the Euclidean distance between the cell centres of two neighbouring cells is 1). $d_H$ is hereafter named the *El Alto distance*, inspired by the peculiar hexagonally shaped urbanisation located in El Alto, Bolivia (Figure S3[1]).

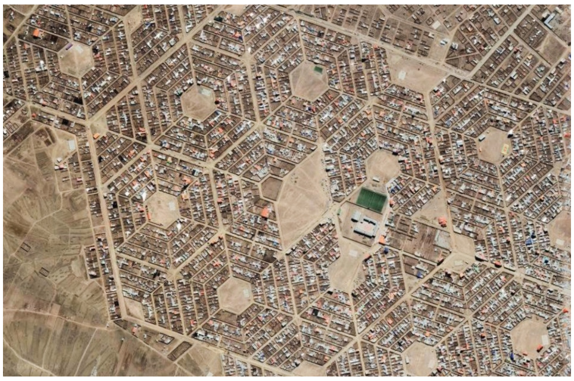

**Figure S3. Hexagonal layout on a neighbourhood of El Alto, Bolivia.**

We recall that we assume signalling between a cell and any of its non-immediate neighbours occurs if it is within reach of protrusions. That is, if the Euclidean distance between both cell centres, $d_E(i,j)$, is less than a certain fixed threshold. It is then possible to write an equivalent definition of $R_k$ using the Euclidean distance by defining the threshold $k-1 < d_E(i,j) \leq k$. However, this is only valid up to a specific value of $k$, so there is an important geometric difference between distances $d_H$ and $d_E$ on the hexagonal lattice.

In other words, above a specific $k$, the distance thresholds define different sets of cells. This first equivalence break occurs when the Euclidean distance to a cell in $R_{k+1}$ is less than the El Alto distance to a cell in $R_k$. Hence, we want to find the smallest integer $k$ such that

$$d_E(i, j \in R_{k+1}) < d_H(i, j \in R_k) \iff d_E(i, j \in R_{k+1}) < k. \tag{S99}$$

To determine the expression for $d_E(i, j \in R_{k+1})$, we first notice that such cells will be positioned towards the middle vertical section of the hexagonal lattice depicted in Figure S4a. To find the first $k$ for which equivalence breaks, two minimal paths are possible, depending on the parity of $k$. The triangles in Figure S4b depict both the shortest Euclidean distance and a corresponding possible El Alto path for each $R_k$. Hence, we have two possibilities for the Euclidean distance, given by

$$d_E(i, j \in R_k) = \begin{cases} \frac{k\sqrt{3}}{2} & k \text{ even}, \\ \frac{\sqrt{3k^2+1}}{2} & k \text{ odd}. \end{cases} \tag{S100}$$

[1]Attribution information: Google Maps, Imagery ©2022 CNES / Airbus, Maxar Technologies, Map data ©2022.

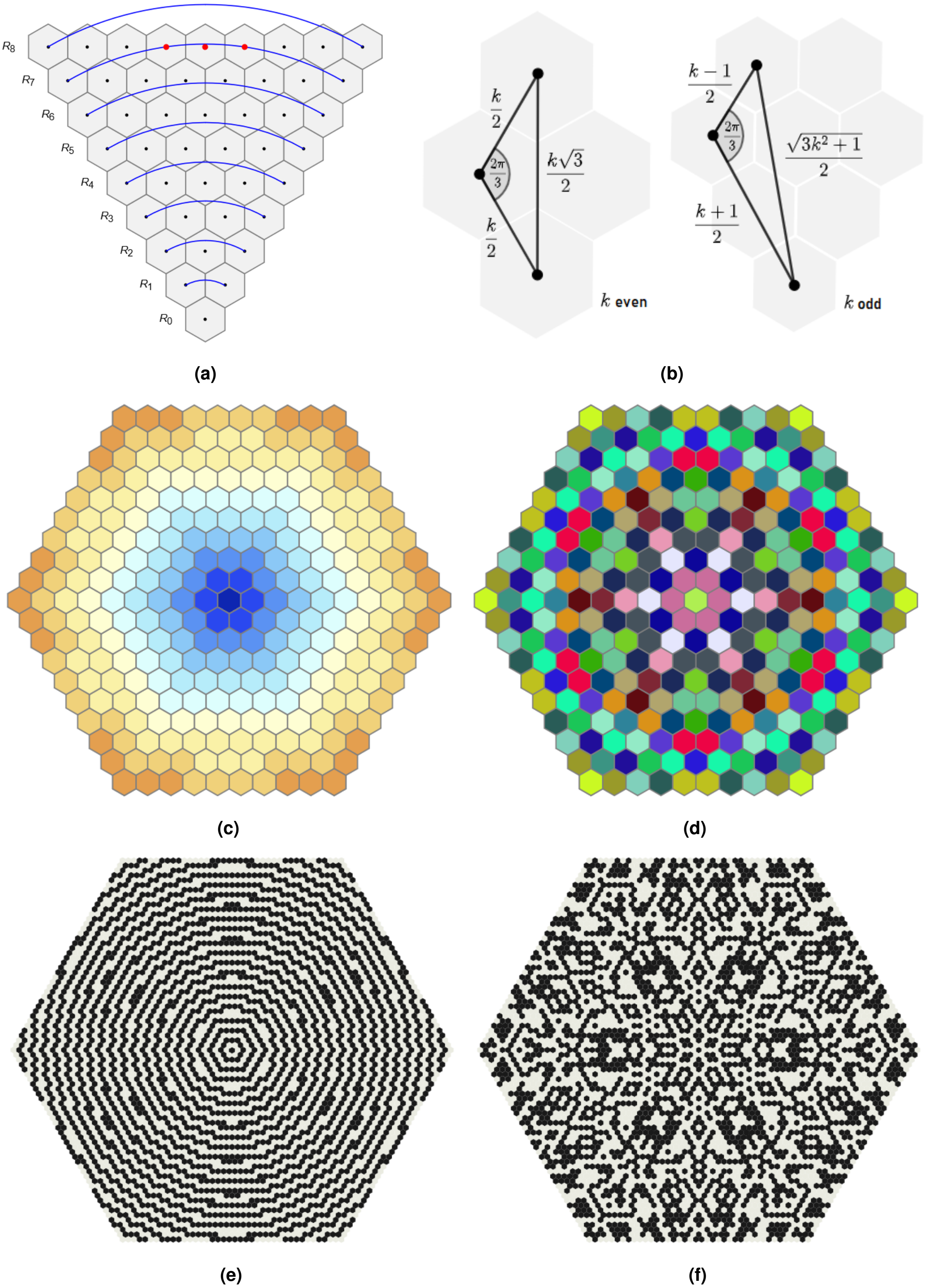


**Figure S4.** (Caption overleaf.)

**Figure S4.** (Overleaf.) **Euclidean and Manhattan distances on hexagonal lattices.**
**(a)** The fixed Euclidean distance (radius of the blue arcs) defined at $R_7$ is greater or equal to the distance of three cell centres (in red) in $R_8$, breaking the definition equivalence. **(b)** Two minimal paths are possible, depending on the parity of $k$. **(c-d)** Signalling rings based on two different distance-based definitions. Cells are in the same ring (same colour) if **(c)** $k-1 < d_E(i,j) \leq k$, or **(d)** they have exactly the same Euclidean distance to cell $i$. $k \leq 10$. **(e-f)** Intricate patterns emerge by alternating colouring of successive signalling rings based on the respective two definitions. $k \leq 40$.

Then, it follows that

$$d_E(i,j \in R_{k+1}) < k \iff \begin{cases} \frac{\sqrt{3(k+1)^2+1}}{2} < k & k \text{ even,} \\ \frac{(k+1)\sqrt{3}}{2} < k & k \text{ odd.} \end{cases} \quad \textbf{(S101)}$$

$$\iff \begin{cases} k > 3+\sqrt{13} \simeq 6.6056 & k \text{ even,} \\ k > \frac{\sqrt{3}}{2-\sqrt{3}} \simeq 6.4641 & k \text{ odd.} \end{cases} \quad \textbf{(S102)}$$

Hence, the equivalence between definitions fails for $k \geq 7$, meaning the $R_k$ Euclidean definition for $R_7$ would already include cells in $R_8$. This observation shows that these two definitions would yield slightly different analysis for signalling at distant rings. We argue, however, that realistic protrusions do not usually reach $R_7$, and therefore the equivalent definitions work for most applications. However, the main text also discusses the analysis of such systems for theoretically longer protrusions under the El Alto distance definition.
Alternative definitions of $R_k$ are possible with fixed Euclidean distance thresholds and can lead to different levels of complexity (Figures S4c-S4f). Interesting patterns emerge when considering signalling rings purely defined by their Euclidean radius. That is, cells belong to the same ring provided they have precisely the same Euclidean distance to the focal cell $i$ (Figure S4d). By alternating the colouring of successive signalling rings, based on this definition and ordered from the distance to $R_0$, somewhat chaotic patterns emerge from such simple labelling (Figure S4f). In other words, we colour cells black or white dependent on their distance from the focal cell, alternating colour whenever a new set of cell centres intersects a circumference with an increasing radius, centered at $R_0$. If the number of greyscale colouring steps is increased (starting with the binary black or white), further fractal-like patterns arise (Video SV4). The only coinciding rings under all definitions are $R_0$ and $R_1$.
Since realistic cells are not regular hexagons, and protrusions are also subject to stochastic effects (such as different lengths and lifespans), we argue that the hexagonally isotropic signalling rings $R_k$ provide a more realistic approximation of the signalling reach of protrusions when compared to the alternative discrete distance-based definition we presented.

### Robustness and pattern selection

As discussed in the main text, Fourier analysis helps in describing pattern selection under LSA. We recall that the solution of the linearised problem is given by

$$\begin{pmatrix} \tilde{n}_{j,k}(t) \\ \tilde{d}_{j,k}(t) \end{pmatrix} = \sum_{q=1}^{N} \sum_{p=1}^{M} \left[ C^{+}_{\bar{q},\bar{p}} e^{\lambda^{+}_{\bar{q},\bar{p}} t} \mathbf{v}^{+}_{\bar{q},\bar{p}} + C^{-}_{\bar{q},\bar{p}} e^{\lambda^{-}_{\bar{q},\bar{p}} t} \mathbf{v}^{-}_{\bar{q},\bar{p}} \right] e^{2\pi \mathrm{i}(\bar{q}j+\bar{p}k)}, \quad \textbf{(S103)}$$

where $\mathbf{v}^{\pm}_{\bar{q},\bar{p}}$ and $\lambda^{\pm}_{\bar{q},\bar{p}}$ are the eigenvectors and eigenvalues associated to $(\bar{q},\bar{p})$, respectively. $C^{+}_{\bar{q},\bar{p}}$ and $C^{-}_{\bar{q},\bar{p}}$ are constants depending on the initial conditions of the problem. Figure S5 shows the comparison between the final patterns from Figure 9 with an SOP cell filtering based on a specific threshold (percentage of steady state solution) and a numerical simulation. Figure S6 shows the dependence of cell fates on the choice of $C^{+}_{\bar{q},\bar{p}}$, as predicted by LSA.
Note also that the bifurcation of $\Omega_{\bar{q},\bar{p}}$ at $\epsilon = 0.4$ (Figure 2b) can be mathematically shown by solving

$$\Omega_{\frac{1}{3},\frac{1}{3}}(\epsilon) = \Omega_{\bar{q},\bar{p}}(\epsilon) \quad \textbf{(S104)}$$

for $\epsilon$ and a minimising pair $(\bar{q},\bar{p}) \notin \{(1/3,1/3),(2/3,2/3)\}$. Given the minimisers in Figure 9b, we have that, with $(\bar{q},\bar{p}) = (1/4,1/4)$ for example,

$$\Omega_{\frac{1}{3},\frac{1}{3}}(\epsilon) = \Omega_{\frac{1}{4},\frac{1}{4}}(\epsilon) \Leftrightarrow \epsilon = 0.4. \quad \textbf{(S105)}$$

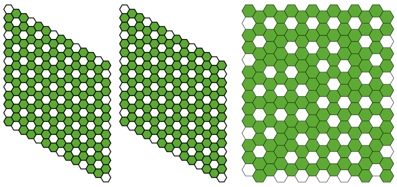

**(a)** $\epsilon = 0$, $d_T = 0$

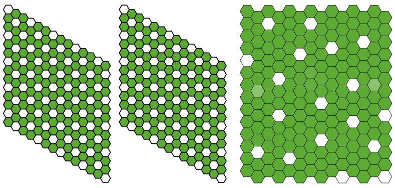

**(b)** $\epsilon = 0.2$, $d_T = 0$

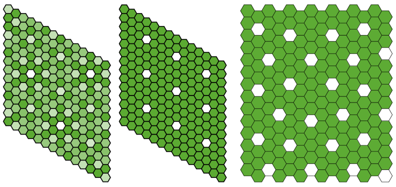

**(c)** $\epsilon = 0.4$, $d_T = 0.64$

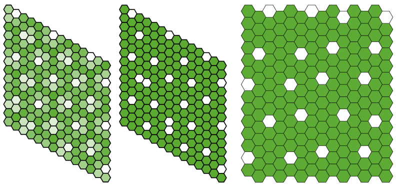

**(d)** $\epsilon = 0.6$, $d_T = 0.73$

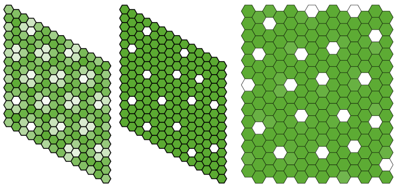

**(e)** $\epsilon = 0.8$, $d_T = 0.82$

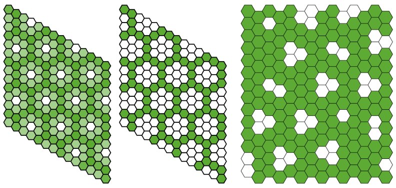

**(f)** $\epsilon = 1$, $d_T = 0.44$

**Figure S5. Pattern selection: threshold solutions and numerical comparisons.**
Here are shown the final patterns predicted by LSA (left), threshold-based SOP cell detection with $d_i > d_T$ (middle) and a numerical simulation (right), for different values of $\epsilon$ and $d_T$. $C^{+}_{\bar{q},\bar{p}} = 1\ \forall(\bar{q},\bar{p})$, $h = k = 4$, $a = 0.01$, $b = 100$ and $\nu = 1$.

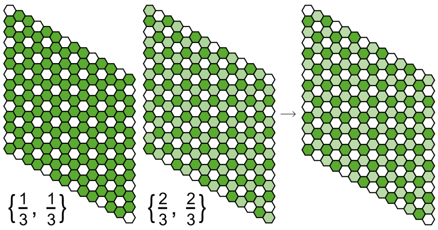


**(a)** $0 \leq \epsilon < 0.4$

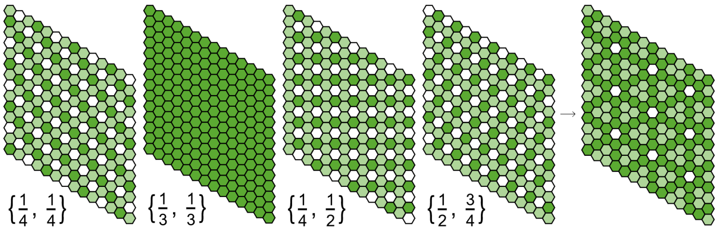


**(b)** $\epsilon = 0.4$

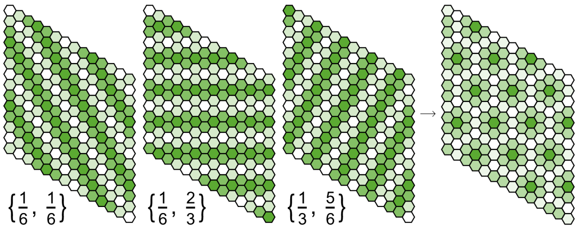


**(c)** $\epsilon = 1$

**Figure S6. Pattern selection: dependence on initial conditions.**
**(a)** $\epsilon = 0$. $C^{+}_{\frac{1}{3},\frac{1}{3}} = 1$, $C^{+}_{\frac{2}{3},\frac{2}{3}} = 0.7\mathrm{i}$. **(b)** $\epsilon = 0.4$, $C^{+}_{\frac{1}{3},\frac{1}{3}} = C^{+}_{\frac{2}{3},\frac{2}{3}} = 0$, $C^{+}_{\bar{q},\bar{p}} = 1$, $\forall \bar{q},\bar{p} \notin \{\{\frac{1}{3},\frac{1}{3}\}\{\frac{2}{3},\frac{2}{3}\}\}$. **(c)** $\epsilon = 1$, $C^{+}_{\bar{q},\bar{p}} = -1$, $\forall \bar{q},\bar{p}$.

### Supplementary Note 2: Weakly nonlinear stability analysis

As discussed in the main text, LSA has failed to predict sparse pattern formation for many values of $\epsilon$. Motivated by theory from reaction-diffusion systems (Turing, 1952; Wollkind et al., 1994), we perform weakly nonlinear stability analysis (WNSA) to better understand the effects of nonlinear terms, based on the amplitude dynamics of the Fourier components corresponding to the fastest growing modes, expanding on LSA to deduce further patterning features derived from long-range signalling. The following discussion is presented in gradually increasing levels of complexity of differential systems.

#### One-dimensional systems

When performing LSA, we have looked for solutions to the following system

$$\frac{du}{dt} = F(u) \tag{S106}$$

of the type

$$u(t) = u^* + \tilde{u}, \tag{S107}$$

where $u^*$ is the homogeneous solution and $\tilde{u}$ is a small perturbation which is solution to the linearised problem $\frac{d\tilde{u}}{dt} = F'(u^*)\tilde{u}$.

Motivated by the theory in reaction-diffusion systems, where the governing equations are nonlinear partial differential equations (PDEs), an approximation to the solution is obtained by introducing the finite amplitude function $A(t,x)$, where $x$ is the spatial variable (Wollkind and Segel, 1970; Wollkind et al., 1984; Wollkind and Vislocky, 1990). For the space-independent system, we take solutions of the type

$$u(t) = \sum_{m=0}^{\infty} v_m A^m(t). \tag{S108}$$

$A$ satisfies the following Landau equation (Aranson and Kramer, 2002)

$$\frac{dA}{dt}(t) = \sum_{m=0}^{\infty} b_m A^m(t), \tag{S109}$$

where $\{v_m\}$ are constants and $\{b_m\}$ are coefficients depending on the problem itself. $\{b_m\}$ are also known as Landau constants. This formulation, often named the Stuart-Watson nonlinear extension (Stuart, 1960; Watson, 1960), allows us to modify the linear expansion approach inherent to Eq. (S107) so that it will be applicable to higher-order terms. Approximate solutions are obtained by a severe truncation of the infinite series. Here, we take

$$u(t) = \sum_{m=0}^{K} v_m A^m(t) + O(A^{K+1}) \tag{S110}$$

and

$$\frac{dA}{dt}(t) = \sum_{m=0}^{K} b_m A^m(t) + O(A^{K+1}). \tag{S111}$$

In this case, substituting into Eq. (S106) leads to

$$\frac{du}{dt} = \sum_{m=0}^{K} v_m \frac{d}{dt}[A^m(t)] \tag{S112}$$

$$= \sum_{m=0}^{K} m v_m A^{m-1}(t) \frac{dA}{dt}(t) \tag{S113}$$

$$= \sum_{m=0}^{K} \sum_{j=0}^{K} m v_m b_j A^{m+j-1}(t). \tag{S114}$$

Solving Eq. (S110)-Eq. (S111) with fixed $K$ for coefficients $\{v_m\}$ and $\{b_m\}$ leads to a sequence of $m$ vector systems, each corresponding to a nonzero term of the form $v_m A^m(t)$ appearing explicitly in Eq. (S110). In general, we start by solving Eq. (S111) and then deduce $\{v_m\}$ and $\{b_m\}$ by differentiating Eq. (S110) and Taylor-expanding $F_A(A) \equiv F(u)$ around $A=0$. Previous works have solved this with $K=3$ for reaction-diffusion equations in different applications (Stephenson and Wollkind, 1995; Bozzini et al., 2015; Liu et al., 2018). We then need to solve the following system

$$\frac{du}{dt} = F_A(A) \tag{S115}$$

for the tuples $(b_0, b_1, b_2, b_3)$ and $(v_0, v_1, v_2, v_3)$, where

$$u(t) = v_0 + v_1 A(t) + v_2 A^2(t) + v_3 A^3(t) + O(A^4) \tag{S116}$$
$$\frac{dA}{dt}(t) = b_0 + b_1 A(t) + b_2 A^2(t) + b_3 A^3(t) + O(A^4). \tag{S117}$$

We take $v_0 \equiv u^*$. Then, substituting Eq. (S116)-Eq. (S117) into $\frac{du}{dt}$ yields

$$\begin{aligned}\frac{du}{dt} &= b_0 v_1 + (b_1 v_1 + 2b_0 v_2) A(t) + (b_2 v_1 + 2b_1 v_2 + 3b_0 v_3) A^2(t) \\ &\quad + (b_3 v_1 + 2b_2 v_2 + 3b_1 v_3) A^3(t) + O(A^4).\end{aligned} \tag{S118}$$

Taylor-expanding $F_A(A)$ around $A=0$ leads to

$$F_A(A) \equiv F(u^* + v_1 A(t) + v_2 A^2(t) + v_3 A^3(t)) \tag{S119}$$
$$\begin{aligned}&\simeq F(u^*) + v_1 F'(u^*) A(t) + \left(\frac{1}{2} v_1^2 F''(u^*) + v_2 F'(u^*)\right) A^2(t) \\ &\quad + \left(\frac{1}{6} v_1^3 F^{(3)}(u^*) + v_2 v_1 F''(u^*) + v_3 F'(u^*)\right) A^3(t) + O(A^4).\end{aligned} \tag{S120}$$

Finally, equating the coefficients of $A^m(t)$ ($0 \le m \le 3$) defines the system of $4$ equations and $7$ variables (excluding the trivial case $v_0 = u^*$)

$$m=0: \quad b_0 v_1 = F(u^*) = 0 \tag{S121}$$
$$m=1: \quad b_1 v_1 + 2b_0 v_2 = v_1 F'(u^*) \tag{S122}$$
$$m=2: \quad b_2 v_1 + 2b_1 v_2 + 3b_0 v_3 = \frac{1}{2} v_1^2 F''(u^*) + v_2 F'(u^*) \tag{S123}$$
$$m=3: \quad b_3 v_1 + 2b_2 v_2 + 3b_1 v_3 = \frac{1}{6} v_1^3 F^{(3)}(u^*) + v_2 v_1 F''(u^*) + v_3 F'(u^*) \tag{S124}$$

from which, with $v_1 \neq 0$, we deduce

$$b_0 = 0 \tag{S125}$$
$$b_1 = F'(u^*) \tag{S126}$$
$$b_2 = \frac{v_1^2 F''(u^*) - 2v_2 F'(u^*)}{2v_1} \tag{S127}$$
$$b_3 = \frac{v_1^4 F^{(3)}(u^*) - 12 v_1 v_3 F'(u^*) + 12 v_2^2 F'(u^*)}{6 v_1^2}. \tag{S128}$$

This method provides a straightforward tool to explicitly determine weakly nonlinear approximations of the form Eq. (S116) for autonomous systems in one dimension.
Amplitude-based solutions of the form Eq. (S116) are assumed to capture the effects of harmonics of the fastest growing modes, appearing as space-independent components. In many cases, solutions have been restricted to amplitude equations of the form

$$\frac{dA}{dt}(t) = \lambda^* A(t) - \kappa A^3(t) + O(A^5), \tag{S129}$$

where $\lambda^*$ is the fastest growth rate of weakly nonlinear perturbations. The sign of the Landau constant $\kappa$ in this differential equation is relevant: if it is positive, then the effect of the term $-\kappa A^3(t)$ is to arrest the exponential growth of $A(t)$ at the value $\sqrt{\lambda^*/\kappa}$ (Turing, 1952). Given $(b_0, b_1, b_2, b_3) = (0, \lambda^*, 0, -\kappa)$, we have that $F'(u^*) = \lambda^*$ and

$$(v_0, v_1, v_2, v_3) = \left( u^*, v_1, \frac{v_1^2 F''(u^*)}{2\lambda^*}, \frac{v_1 \left( 6\kappa\lambda^* + \lambda^* v_1^2 F^{(3)}(u^*) + 3 v_1^2 F''(u^*)^2 \right)}{12{\lambda^*}^2} \right) \quad \textbf{(S130)}$$

by Eq. (S125)-Eq. (S128) and where $v_1$ is arbitrary. Without loss of generality, we take $v_1 = 1$. Hence, we are interested in solutions of the form

$$u(t) = u^* + A(t) + \frac{F''(u^*)}{2\lambda^*} A^2(t) + \frac{\left( 6\kappa\lambda^* + \lambda^* F^{(3)}(u^*) + 3F''(u^*)^2 \right)}{12{\lambda^*}^2} A^3(t). \quad \textbf{(S131)}$$

$A(t)$ is a solution of Eq. (S117), given by

$$A(t) = \frac{\sqrt{\lambda^*} e^{\lambda(t-C_1)}}{\sqrt{\kappa e^{2\lambda^*(t-C_1)} - 1}}, \quad \textbf{(S132)}$$

where $C_1$ is an integration constant and $\kappa > e^{-2\lambda^*(t-C_1)}$. We shall return to this solution, after discussing the multidimensional analysis next.

#### Multidimensional systems

The generalisation of WNSA to multidimensional systems requires further machinery from calculus. Before we delve into the Notch-motivated two-dimensional system, we present the general Taylor expansion of vector-valued multivariable functions.

Consider a function $\mathbf{F} : \mathbb{R}^n \to \mathbb{R}^m$ given by $\mathbf{F}(\mathbf{u}) = (F_1(\mathbf{u}), ..., F_m(\mathbf{u}))$, where $\mathbf{u} = (u_0, ..., u_n)$. The general $K$th-order Taylor expansion of $\mathbf{F}(\mathbf{u} + \mathbf{u}_0)$ about a point $\mathbf{u}_0 \in \mathbb{R}^n$ is given by

$$\mathbf{F}(\mathbf{u} + \mathbf{u}_0) \simeq \sum_{j=0}^{K} \frac{1}{j!} (D^j \mathbf{F})_{\mathbf{u}_0} [(\mathbf{u})^j], \quad \textbf{(S133)}$$

where the Frechet-derivative terms $(D^j \mathbf{F})_{\mathbf{u}_0}[(\mathbf{u})^j]$ may be written in vector form as

$$(D^j \mathbf{F})_{\mathbf{u}_0}[(\mathbf{u})^j] = \begin{pmatrix} \sum_{i_1,\ldots,i_j=1}^{n} \frac{\partial^j F_1}{\partial u_{i_1} \cdots \partial u_{i_j}} (\mathbf{u}_0)(u_{i_1} \cdots u_{i_j}) \\ \vdots \\ \sum_{i_1,\ldots,i_j=1}^{n} \frac{\partial^j F_m}{\partial u_{i_1} \cdots \partial u_{i_j}} (\mathbf{u}_0)(u_{i_1} \cdots u_{i_j}) \end{pmatrix} \quad \textbf{(S134)}$$

using the notation

$$\sum_{i_1,\ldots,i_j=1}^{n} = \sum_{i_1=1}^{n} \cdots \sum_{i_j=1}^{n}. \quad \textbf{(S135)}$$

For a two-variable two-dimensional ($n = m = 2$) vector function $\mathbf{F}(\mathbf{u}) = (F_1(\mathbf{u}), F_2(\mathbf{u}))$, where $\mathbf{u} = (u_1, u_2)$, the third-order Taylor expansion of such function around $\mathbf{u}_0 = \mathbf{u}^* = (u_1^*, u_2^*)$ is given by

$$\begin{aligned} \mathbf{F}(\mathbf{u}) \simeq & (D^0 \mathbf{F})_{\mathbf{u}^*}[(\mathbf{u} - \mathbf{u}^*)^0] + (D^1 \mathbf{F})_{\mathbf{u}^*}[(\mathbf{u} - \mathbf{u}^*)^1] \\ & + \frac{1}{2} (D^2 \mathbf{F})_{\mathbf{u}^*}[(\mathbf{u} - \mathbf{u}^*)^2] + \frac{1}{6} (D^3 \mathbf{F})_{\mathbf{u}^*}[(\mathbf{u} - \mathbf{u}^*)^3], \end{aligned} \quad \textbf{(S136)}$$

which may be simplified as follows

$$(D^0 \mathbf{F})_{\mathbf{u}^*}[(\mathbf{u} - \mathbf{u}^*)^0] = \mathbf{F}(\mathbf{u}^*) \quad \textbf{(S137)}$$

$$(D^1 \mathbf{F})_{\mathbf{u}^*}[(\mathbf{u} - \mathbf{u}^*)^1] = \begin{pmatrix} \sum_{i_1=1}^{2} \frac{\partial F_1}{\partial u_{i_1}} (\mathbf{u}^*) \left[ (u_{i_1} - u_{i_1}^*) \right] \\ \sum_{i_1=1}^{2} \frac{\partial F_2}{\partial u_{i_1}} (\mathbf{u}^*) \left[ (u_{i_1} - u_{i_1}^*) \right] \end{pmatrix} = \mathbf{J}(\mathbf{u}^*)(\mathbf{u} - \mathbf{u}^*) \quad \textbf{(S138)}$$

$$(D^2 \mathbf{F})_{\mathbf{u}^*}[(\mathbf{u} - \mathbf{u}^*)^2] = \begin{pmatrix} \sum_{i_1=1}^{2} \sum_{i_2=1}^{2} \frac{\partial^2 F_1}{\partial u_{i_1} \partial u_{i_2}} (\mathbf{u}^*) \left[ (u_{i_1} - u_{i_1}^*)(u_{i_2} - u_{i_2}^*) \right] \\ \sum_{i_1=1}^{2} \sum_{i_2=1}^{2} \frac{\partial^2 F_2}{\partial u_{i_1} \partial u_{i_2}} (\mathbf{u}^*) \left[ (u_{i_1} - u_{i_1}^*)(u_{i_2} - u_{i_2}^*) \right] \end{pmatrix} \quad \textbf{(S139)}$$

$$= \begin{pmatrix} (\mathbf{u}-\mathbf{u}^*)^T \mathbf{H}_1(\mathbf{u}^*)(\mathbf{u}-\mathbf{u}^*) \\ (\mathbf{u}-\mathbf{u}^*)^T \mathbf{H}_2(\mathbf{u}^*)(\mathbf{u}-\mathbf{u}^*) \end{pmatrix} \tag{S140}$$

$$(D^3\mathbf{F})_{\mathbf{u}^*}[(\mathbf{u}-\mathbf{u}^*)^3]$$
$$= \begin{pmatrix} \sum_{i_1=1}^2 \sum_{i_2=1}^2 \sum_{i_3=1}^2 \frac{\partial^3 F_1}{\partial u_{i_1} \partial u_{i_2} \partial u_{i_3}}(\mathbf{u}^*) \left[ (u_{i_1}-u_{i_1}^*)(u_{i_2}-u_{i_2}^*)(u_{i_3}-u_{i_3}^*) \right] \\ \sum_{i_1=1}^2 \sum_{i_2=1}^2 \sum_{i_3=1}^2 \frac{\partial^3 F_2}{\partial u_{i_1} \partial u_{i_2} \partial u_{i_3}}(\mathbf{u}^*) \left[ (u_{i_1}-u_{i_1}^*)(u_{i_2}-u_{i_2}^*)(u_{i_3}-u_{i_3}^*) \right] \end{pmatrix}. \tag{S141}$$

$\mathbf{J}$ is the Jacobian matrix of $\mathbf{F}$, and $\mathbf{H}_1$ and $\mathbf{H}_2$ are the Hessian matrices of functions $F_1$ and $F_2$, respectively. The third derivative term given by $(D^3\mathbf{F})_{\mathbf{u}^*}[(\mathbf{u}-\mathbf{u}^*)^3]$ has no straightforward simplification. Hence

$$\mathbf{F}(\mathbf{u}) \simeq \mathbf{F}(\mathbf{u}^*) + \mathbf{J}(\mathbf{u}^*)(\mathbf{u}-\mathbf{u}^*)$$
$$+ \frac{1}{2} \begin{pmatrix} (\mathbf{u}-\mathbf{u}^*)^T \mathbf{H}_1(\mathbf{u}^*)(\mathbf{u}-\mathbf{u}^*) \\ (\mathbf{u}-\mathbf{u}^*)^T \mathbf{H}_2(\mathbf{u}^*)(\mathbf{u}-\mathbf{u}^*) \end{pmatrix} + \frac{1}{6}(D^3\mathbf{F})_{\mathbf{u}^*}[(\mathbf{u}-\mathbf{u}^*)^3]. \tag{S142}$$

In the following, we drop the differential evaluation at the homogeneous state $\mathbf{u}^*$ when convenient. Again, we fix $K=3$ and consider solutions to

$$\frac{d\mathbf{u}}{dt} = \mathbf{F}(\mathbf{u}) \tag{S143}$$

of the form

$$\mathbf{u}(t) = \mathbf{v}_0 + \mathbf{v}_1 A(t) + \mathbf{v}_2 A^2(t) + \mathbf{v}_3 A^3(t) + O(A^4) \tag{S144}$$
$$= \begin{pmatrix} v_{01} \\ v_{02} \end{pmatrix} + \begin{pmatrix} v_{11} \\ v_{12} \end{pmatrix} A(t) + \begin{pmatrix} v_{21} \\ v_{22} \end{pmatrix} A^2(t) + \begin{pmatrix} v_{31} \\ v_{32} \end{pmatrix} A^3(t) + O(A^4), \tag{S145}$$

where $A(t)$ satisfies Eq. (S117) and $\mathbf{v}_m$ ($0 \le m \le 3$) are now constant vectors. Using the general Taylor expansion Eq. (S142), we now aim to find the tuples $(b_0, b_1, b_2, b_3)$, $(v_{01}, v_{11}, v_{21}, v_{31})$ and $(v_{02}, v_{12}, v_{22}, v_{32})$ such that the weakly nonlinear approximation holds in the two-dimensional system.
Substituting Eq. (S145) into $\frac{d\mathbf{u}}{dt}$ we get, for $j \in \{1,2\}$,

$$\frac{du_j}{dt} = b_0 v_{1,j} + (b_1 v_{1,j} + 2b_0 v_{2,j}) A(t) + (b_2 v_{1,j} + 2b_1 v_{2,j} + 3b_0 v_{3,j}) A^2(t)$$
$$+ (b_3 v_{1,j} + 2b_2 v_{2,j} + 3b_1 v_{3,j}) A^3(t) + O(A^4). \tag{S146}$$

Given Eq. (S142), we have that the Taylor expansion of $\mathbf{F}_A(A) \equiv \mathbf{F}(\mathbf{u})$ around $A=0$ satisfies

$$F_{Aj}(A) = F_j(\mathbf{v}_1 A(t) + \mathbf{v}_2 A^2(t) + \mathbf{v}_3 A^3(t)) \tag{S147}$$
$$\simeq \mathbf{T}_{0,j}(\mathbf{v}) + \mathbf{T}_{1,j}(\mathbf{v}) A(t) + \mathbf{T}_{2,j}(\mathbf{v}) A^2(t) + \mathbf{T}_{3,j}(\mathbf{v}) A^3(t) + O(A^4), \tag{S148}$$

where $\mathbf{v} \equiv (\mathbf{v}_0, \mathbf{v}_1, \mathbf{v}_2, \mathbf{v}_3)$ and

$$\mathbf{T}_{0,j}(\mathbf{v}) = F_j = 0 \tag{S149}$$
$$\mathbf{T}_{1,j}(\mathbf{v}) = v_{11} \frac{\partial F_j}{\partial u_1} + v_{12} \frac{\partial F_j}{\partial u_2} \tag{S150}$$
$$\mathbf{T}_{2,j}(\mathbf{v}) = v_{21} \frac{\partial F_j}{\partial u_1} + v_{22} \frac{\partial F_j}{\partial u_2} + \frac{v_{11}^2}{2} \frac{\partial^2 F_j}{\partial u_1^2} + \frac{v_{12}^2}{2} \frac{\partial^2 F_j}{\partial u_2^2} + v_{11} v_{12} \frac{\partial^2 F_j}{\partial u_1 \partial u_2} \tag{S151}$$
$$\mathbf{T}_{3,j}(\mathbf{v}) = v_{31} \frac{\partial F_j}{\partial u_1} + v_{32} \frac{\partial F_j}{\partial u_2}$$
$$+ v_{11} v_{21} \frac{\partial^2 F_j}{\partial u_1^2} + v_{12} v_{22} \frac{\partial^2 F_j}{\partial u_2^2} + (v_{21} v_{12} + v_{11} v_{22}) \frac{\partial^2 F_j}{\partial u_1 \partial u_2}$$
$$+ \frac{v_{11}^3}{6} \frac{\partial^3 F_j}{\partial u_1^3} + \frac{v_{12}^3}{6} \frac{\partial^3 F_j}{\partial u_2^3} + \frac{v_{11}^2 v_{12}}{2} \frac{\partial^3 F_j}{\partial u_1^2 \partial u_2} + \frac{v_{11} v_{12}^2}{2} \frac{\partial^3 F_j}{\partial u_1 \partial u_2^2}. \tag{S152}$$

Equating the coefficients of Eq. (S146) and Eq. (S148) yields an intricate system of 8 equations and 10 variables (excluding the trivial cases $v_{01} = u_1^*$ and $v_{02} = u_2^*$), leading to a series of relations between $b_m$ and $v_{m',j}$ ($1 \le m, m' \le 3$, $j \in \{1,2\}$). By defining $\mathbf{T}_m \equiv (\mathbf{T}_{m,1}, \mathbf{T}_{m,2})^T$, for $0 \le m \le K$, we have

$$\mathbf{T}_0(\mathbf{v}) = \mathbf{0} \tag{S153}$$

$$\mathbf{T}_1(\mathbf{v}) = \mathbf{J}(\mathbf{u}^*)\mathbf{v}_1 \tag{S154}$$

$$\mathbf{T}_2(\mathbf{v}) = \mathbf{J}(\mathbf{u}^*)\mathbf{v}_2 + \frac{1}{2}\begin{pmatrix}\mathbf{v}_1^T\mathbf{H}_1(\mathbf{u}^*)\mathbf{v}_1\\ \mathbf{v}_1^T\mathbf{H}_2(\mathbf{u}^*)\mathbf{v}_1\end{pmatrix} \tag{S155}$$

$$\mathbf{T}_3(\mathbf{v}) = \mathbf{J}(\mathbf{u}^*)\mathbf{v}_3 + \frac{1}{2}\begin{pmatrix}\mathbf{v}_2^T\mathbf{H}_1(\mathbf{u}^*)\mathbf{v}_1\\ \mathbf{v}_2^T\mathbf{H}_2(\mathbf{u}^*)\mathbf{v}_1\end{pmatrix} + \frac{1}{6}(D^3\mathbf{F})_{\mathbf{u}^*}[(\mathbf{v}_1)^3] \tag{S156}$$

and so the coefficient relations are given by

$$b_0\mathbf{v}_1 = \mathbf{0} \tag{S157}$$

$$b_1\mathbf{v}_1 = \mathbf{J}(\mathbf{u}^*)\mathbf{v}_1 \tag{S158}$$

$$(2b_1\mathbf{I} - \mathbf{J}(\mathbf{u}^*))\mathbf{v}_2 = \frac{1}{2}\begin{pmatrix}\mathbf{v}_1^T\mathbf{H}_1(\mathbf{u}^*)\mathbf{v}_1\\ \mathbf{v}_1^T\mathbf{H}_2(\mathbf{u}^*)\mathbf{v}_1\end{pmatrix} - b_2\mathbf{v}_1 \tag{S159}$$

$$(3b_1\mathbf{I} - \mathbf{J}(\mathbf{u}^*))\mathbf{v}_3 = \frac{1}{2}\begin{pmatrix}\mathbf{v}_2^T\mathbf{H}_1(\mathbf{u}^*)\mathbf{v}_1\\ \mathbf{v}_2^T\mathbf{H}_2(\mathbf{u}^*)\mathbf{v}_1\end{pmatrix} + \frac{1}{6}(D^3\mathbf{F})_{\mathbf{u}^*}[(\mathbf{v}_1)^3] - b_3\mathbf{v}_1 - 2b_2\mathbf{v}_2. \tag{S160}$$

We now motivate a choice for $\mathbf{v}_1 = (v_{11}, v_{12})$. Notice first that this method mimics the LSA approach when $K = 1$, $\mathbf{v}_0 = \mathbf{u}^*$ and $b_0 = 0$. Eq. (S158) holds provided $b_1$ and $\mathbf{v}_1$ are an eigenvalue and a corresponding eigenvector of the Jacobian matrix $\mathbf{J}$ of $\mathbf{F}$ evaluated at $\mathbf{u}^*$. Assuming $A(t) = e^{\lambda^* t}$, we take $b_1 = \lambda^*$ and $\mathbf{v}_1 = \mathbf{v}^*$, where $\lambda^*$ is the fastest growth rate and $\mathbf{v}^*$ its corresponding eigenvector (which is not unique), as in Eq. (66)-Eq. (67). The general solution would, in this case, correspond to the asymptotic solution in Eq. (68). The remaining coefficient relations between Eq. (S146) and Eq. (S148) follow from this choice of $\mathbf{v}_1$ via the other coefficient identities.

#### WNSA of Notch-Delta signalling dynamics

In the following, we take $\mathbf{u}_i = (n_i, d_i)$ and

$$\frac{d\mathbf{u}_i}{dt} = \mathbf{F}(n_i, d_i) = \begin{pmatrix}F_1(n_i,d_i)\\ F_2(n_i,d_i)\end{pmatrix} = \begin{pmatrix}f(\langle d_i\rangle) - n_i\\ \nu(g(n_i) - d_i)\end{pmatrix} \tag{S161}$$

for each cell $i$. We may interchange cell indexation between $i$ and $(j,k)$ as convenient, as well as the conditions $r \neq i$, $r \neq (j,k)$ and $\Delta jk \in S$ (see previous sections for details on such notation). As previously seen, we aim to extend the linear approach to consider solutions of Eq. (S161) in the harmonic form

$$\mathbf{u}_i(t) = \mathbf{u}^* + \mathbf{v}_1^i A(t) + \mathbf{v}_2^i A^2(t) + \mathbf{v}_3^i A^3(t) + O(A^4), \tag{S162}$$

where

$$\frac{dA}{dt}(t) = \lambda^* A(t) - \kappa A^3(t) + O(A^5). \tag{S163}$$

We have seen that discrete Fourier transforms Eq. (S62)-Eq. (S65) may be used to decouple the original system of $2NM$ equations ($NM$ is the number of cells on a $N \times M$ hexagonal lattice). We recall from Eq. (S61) that, when $K = 1$, linearisation led to

$$\frac{d\mathbf{u}_i}{dt} \simeq \mathbf{J}\mathbf{u}_i + \sum_{r\neq i}\mathbf{J}^r\mathbf{u}_r, \tag{S164}$$

which was then rewritten by introducing the following coupling function derived from the Jacobian matrix (with $i \equiv (j,k)$ in the case of a two-dimensional lattice)

$$\mathbf{\Omega}_{\bar{q},\bar{p}} \equiv \mathbf{\Omega}_{\bar{q},\bar{p}}(q,p) = \sum_{\Delta jk\in S}\mathbf{J}^{\Delta jk}e^{-2\pi \mathrm{i}(\bar{q}\Delta j + \bar{p}\Delta k)} \tag{S165}$$

leading to the decoupled linearised problem

$$\frac{d}{dt}\begin{pmatrix}\xi_{q,p}\\ \eta_{q,p}\end{pmatrix} \simeq \mathbf{L}_{\bar{q},\bar{p}}\begin{pmatrix}\xi_{q,p}\\ \eta_{q,p}\end{pmatrix}, \tag{S166}$$

where $\mathbf{L}_{\bar{q},\bar{p}} = \mathbf{J} + \mathbf{\Omega}_{\bar{q},\bar{p}}$. Given exponential-based solutions of the linearised Eq. (S166), our problem relied then on minimising $\mathbf{\Omega}_{\bar{q},\bar{p}}$ in order to find the fastest growing modes $(\bar{q},\bar{p})$. This problem changes and becomes relatively trickier in the case of WNSA due to multiple mathematical obstacles, as discussed below.
The extension of Eq. (S165) to the weakly nonlinear solution Eq. (S162) is not trivial and, in general, we should not expect a higher-order extension of decomposition Eq. (S164) to occur due to cross-derivative terms. To see this,

take a general function $\mathbf{F}(\mathbf{u}) = (F_1(\mathbf{u}), F_2(\mathbf{u}))$, where $\mathbf{u} = (n_i, d_i, d_{i-1}, d_{i+1})$ (corresponding, for example, to the one-dimensional ring of cells in Example 1.2). The second-order term of the Taylor expansion for $F_1$ around $\mathbf{u}^*$ includes the cross-derivative terms

$$n_i d_{i-1} \frac{\partial^2 F_1}{\partial n_i \partial d_{i-1}} + n_i d_{i+1} \frac{\partial^2 F_1}{\partial n_i \partial d_{i+1}} + d_i d_{i-1} \frac{\partial^2 F_1}{\partial d_i \partial d_{i-1}} + d_i d_{i+1} \frac{\partial^2 F_1}{\partial d_i \partial d_{i+1}}. \quad \textbf{(S167)}$$

Such terms would naturally complicate the decoupling of the relative-index terms $d_{i-1}$ and $d_{i+1}$. However, in our case, with $\mathbf{F}$ given by Eq. (S161), the derivatives in Eq. (S167) are all zero (and consequently any higher-order cross-derivatives). Thus we may hope for a smoother decoupling. Hence, according to Eq. (S133) and generalising Eq. (S164), we may write

$$\frac{d\mathbf{u}_i}{dt} \simeq \sum_{j=0}^{K} \frac{1}{j!} (D^j \mathbf{F})_{\mathbf{u}^*}[(\mathbf{u}_i)^j] + \sum_{r \neq i} \left[ \sum_{j=0}^{K} \frac{1}{j!} (D^j \mathbf{F})_{\mathbf{u}^*}[(\mathbf{u}_r)^j] \right]. \quad \textbf{(S168)}$$

All that remains now is to find a simplification of the right-hand side term of Eq. (S168) so that the decoupling is complete and we may write the entire expression as a function of $(n_i, d_i)$. In other words, we aim to determine the coefficient contribution of the coupling term to Eq. (S157)-Eq. (S160).
To simplify notation and considering the candidate solution Eq. (S162), we begin by Taylor-expanding $\mathbf{F}_A(A) \equiv \mathbf{F}(\mathbf{u})$ around $A = 0$, rewriting Eq. (S168) as follows

$$\mathbf{F}_A(A) \simeq \sum_{m=0}^{3} \mathbf{T}_m^i(\mathbf{v}) A^m(t) + \sum_{r \neq i} \sum_{m=0}^{3} \mathbf{T}_m^r(\mathbf{v}) A^m(t). \quad \textbf{(S169)}$$

With $i \equiv (j,k)$, we apply the following change of variables (Fourier transform)

$$\boldsymbol{\zeta}_{q,p} = \frac{1}{NM} \sum_{k=1}^{M} \sum_{j=1}^{N} \mathbf{u}_{j,k} e^{-2\pi \mathrm{i}(\bar{q}j + \bar{p}k)} \quad \textbf{(S170)}$$

$$= \frac{1}{NM} \sum_{k=1}^{M} \sum_{j=1}^{N} \left[ \mathbf{u}^* + \mathbf{v}_1^{j,k} A(t) + \mathbf{v}_2^{j,k} A^2(t) + \mathbf{v}_3^{j,k} A^3(t) \right] e^{-2\pi \mathrm{i}(\bar{q}j + \bar{p}k)} \quad \textbf{(S171)}$$

$$= \sum_{m=0}^{3} \boldsymbol{\zeta}_{q,p}^m, \quad \textbf{(S172)}$$

where

$$\boldsymbol{\zeta}_{q,p}^m \equiv \frac{1}{NM} \sum_{k=1}^{M} \sum_{j=1}^{N} \mathbf{v}_m^{j,k} A^m(t) e^{-2\pi \mathrm{i}(\bar{q}j + \bar{p}k)}. \quad \textbf{(S173)}$$

From the methods discussed before, it follows that

$$\frac{d\boldsymbol{\zeta}_{q,p}}{dt} = \frac{1}{NM} \sum_{k=1}^{M} \sum_{j=1}^{N} \frac{d\mathbf{u}_{j,k}}{dt} e^{-2\pi \mathrm{i}(\bar{q}j + \bar{p}k)} \quad \textbf{(S174)}$$

$$= \frac{1}{NM} \sum_{k=1}^{M} \sum_{j=1}^{N} \left[ \sum_{m=0}^{3} \mathbf{T}_m^{j,k}(\mathbf{v}) A^m(t) + \sum_{r \neq (j,k)} \sum_{m=0}^{3} \mathbf{T}_m^r(\mathbf{v}) A^m(t) \right] e^{-2\pi \mathrm{i}(\bar{q}j + \bar{p}k)}. \quad \textbf{(S175)}$$

We now aim to decouple the terms

$$\boldsymbol{\Omega}_{\bar{q},\bar{p}}^{3,m}(\mathbf{v}) \equiv \frac{1}{NM} \sum_{k=1}^{M} \sum_{j=1}^{N} \left[ \sum_{r \neq (j,k)} \mathbf{T}_m^r(\mathbf{v}) A^m(t) \right] e^{-2\pi \mathrm{i}(\bar{q}j + \bar{p}k)} \quad \textbf{(S176)}$$

for each $0 \leq m \leq 3$:

- $m = 0$. $\boldsymbol{\Omega}_{\bar{q},\bar{p}}^{3,0}(\mathbf{v}) = \mathbf{0}$.

- $m=1$. This case mimics the deduction of Eq. (S66), as follows

$$\boldsymbol{\Omega}^{3,1}_{\bar{q},\bar{p}}(\mathbf{v}) = \frac{1}{NM}\sum_{k=1}^{M}\sum_{j=1}^{N}\left[\sum_{r\neq(j,k)} \mathbf{T}^r_1(\mathbf{v})A(t)\right] e^{-2\pi\mathrm{i}(\bar{q}j+\bar{p}k)} \tag{S177}$$

$$= \frac{A(t)}{NM}\sum_{k=1}^{M}\sum_{j=1}^{N}\left[\sum_{r\neq(j,k)} \mathbf{J}^r\mathbf{v}^r_1\right] e^{-2\pi\mathrm{i}(\bar{q}j+\bar{p}k)} \tag{S178}$$

$$= \frac{A(t)}{NM}\sum_{\Delta jk\in S} \mathbf{J}^{\Delta jk}\left[\sum_{k=1}^{M}\sum_{j=1}^{N}\mathbf{v}_1^{(j,k)+\Delta jk} e^{-2\pi\mathrm{i}(\bar{q}j+\bar{p}k)}\right] \tag{S179}$$

$$= \sum_{\Delta jk\in S}\mathbf{J}^{\Delta jk}\boldsymbol{\zeta}^1_{q,p}e^{-2\pi\mathrm{i}(\bar{q}\Delta j+\bar{p}\Delta k)} \tag{S180}$$

$$= \boldsymbol{\zeta}^1_{q,p}\boldsymbol{\Omega}_{\bar{q},\bar{p}}, \tag{S181}$$

where $\boldsymbol{\Omega}_{\bar{q},\bar{p}}$ is given by Eq. (S165).

- $m=2$. We have

$$\boldsymbol{\Omega}^{3,2}_{\bar{q},\bar{p}}(\mathbf{v}) = \frac{1}{NM}\sum_{k=1}^{M}\sum_{j=1}^{N}\left[\sum_{r\neq(j,k)} \mathbf{T}^r_2(\mathbf{v})A^2(t)\right] e^{-2\pi\mathrm{i}(\bar{q}j+\bar{p}k)} \tag{S182}$$

$$= \frac{A^2(t)}{NM}\sum_{k=1}^{M}\sum_{j=1}^{N}\left[\sum_{r\neq(j,k)}\left[\mathbf{J}^r\mathbf{v}^r_2+\frac{1}{2}\begin{pmatrix}\mathbf{v^r_1}^T\mathbf{H}^r_1\mathbf{v}^r_1\\ \mathbf{v^r_1}^T\mathbf{H}^r_2\mathbf{v}^r_1\end{pmatrix}\right]\right] e^{-2\pi\mathrm{i}(\bar{q}j+\bar{p}k)}. \tag{S183}$$

While the term involving $\mathbf{J}^r$ can be simplified like the $m=1$ case, the other term yields a higher level of complexity (Remark 2.1). To see this, we exclude both $A^2(t)$ and the index sum, and track the first component of such term, as follows

$$\frac{1}{2NM}\sum_{k=1}^{M}\sum_{j=1}^{N}\left[{v^r_{11}}^2\frac{\partial^2F_1}{\partial {v^r_{11}}^2}+{v^r_{12}}^2\frac{\partial^2F_1}{\partial {v^r_{12}}^2}+2v^r_{11}v^r_{12}\frac{\partial^2F_1}{\partial v^r_{11}\partial v^r_{12}}\right]e^{-2\pi\mathrm{i}(\bar{q}j+\bar{p}k)}. \tag{S184}$$

Given the type of variable change Eq. (S170), we do not expect, in general, to be able to manipulate Eq. (S184) so that it is written in terms of $\boldsymbol{\zeta}^2_{q,p}$ in order to second-order decouple the original system. The same can be argued for the case $m=3$.

- $m=3$. See $m=2$ and discussion below.

Given the complexity generated by the cases $m=2$ and $m=3$ any methodology as systematic as the linear case seems to be out of reach. Therefore, WNSA is insufficient to describe quantitative dynamics of long-range signalling, without further assumptions.

We have presented the main methodology behind a potential framework for weakly nonlinear analysis of translationally invariant Notch-Delta systems. Considering different changes of variables or taking cell-dependent amplitude functions $A_{q,p}(t)$ could help in simplifying decoupling. In the main text, we discuss an additional alternative to our LSA and WNSA approaches for studying Notch-Delta systems, as well as some suggestions for how these methodologies might be improved.

*Remark 2.1* (High-order decoupling) Part of the problem in decoupling the second-order term in Eq. (S182) relies on understanding how a term in the form

$$\sum_{j=1}^{N} a_j^2 e^{\frac{2\pi\mathrm{i}}{N}j} \tag{S185}$$

relates to the quadratic form

$$\left(\sum_{j=1}^{N} a_j e^{\frac{2\pi\mathrm{i}}{N}j}\right)^2, \tag{S186}$$

where $a_j = v^r_{11}$, for example. While a linear manipulation does not seem promising in this case (compared to the $m=1$ case), alternative approaches might hint at further simplification.

### Supplementary Note 3: Simulation parameters

Table 1 shows the model parameters used in the simulations shown in the main text. In all simulations, initial conditions $n_i(0)$ and $d_i(0)$ have arbitrary values between $0$ and $0.1$.

| Signalling | $a$ | $b$ | $h$ | $k$ |
|---|---|---|---|---|
| All figures | 0.01 | 100 | 6 | 6 |

| Figures | $p_\ell$ | $\epsilon$ |
|---|---|---|
| 1c-1f | 2.1 | $\{0, 0.4, 0.6, 1\}$ |
| 2g-2i | 2.1 | $\{0.2, 0.4, 0.6\}$ |
| 3a-3b | 2.1 | $\{0, 0.2, 0.4, 0.6, 0.8, 1\}$ |
| 4a-4d | 2.1 | $\{0.039, 0.85\}$ |
| 5a-5c | 4 | $\{0.2, 0.4, 0.6\}$ |
| 6a-6c | $\{2, 3, 4\}$ | 0.6 |
| 7b-7e | 2.1 | 0.6 |

**Table 1.** Simulation parameters (main text simulations).
